# Local optogenetic control of genome editing and tumorigenesis *in vivo* using wireless implantable optoelectronics

**DOI:** 10.1101/2025.06.12.659214

**Authors:** Trisha Bansal, Min-Kyu Lee, Courtney Coleman, Young-Mee Kim, Yaner Chen, Chad R. Haney, Odile David, Varsha Sreekanth, Raudel Avila, Jacob Matsche, Shaluah Vijeth, Dane Hintermueller, Kento Yamagishi, Jiheon Kang, Katie Mowry, Minsung Kim, Yuru Liu, Peter T. Toth, Steve S-Y. Lee, Sophianne Loh, Sachin Clark, Huong Huynh, Youngdo Kim, Anthony R. Banks, Yonggang Huang, Cameron H. Good, Jalees Rehman, John A. Rogers, Andrei V. Karginov

## Abstract

Precise spatial regulation of site-specific DNA recombination (SSR) *in vivo* remains a challenge due to limited tunability of current platforms. Here, we present an optogenetic approach that overcome these limitations by employing engineered light-regulated recombinase E-LightR-Cre and tunable wireless implantable optoelectronic devices. E-LightR-Cre meets the key criteria for spatial regulation of SSR *in vivo*, showing no detectable activity in the dark, while demonstrating robust activation upon blue-light illumination. To achieve local E-LightR-Cre activation in murine lungs, we developed wireless, fully-implantable optoelectronic devices enabling focal illumination with no discernible organ damage. By modulating illumination intensity and duration, we can control the size of the activated area. Local expression of oncogenic KRas-G12D in a photoactivated subpopulation of cells *in vitro* revealed rapid reprogramming of the mutant expressing cells and their non-activated neighbors. Light-guided activation of E-LightR-Cre in mouse lungs resulted in focal expression of a reporter gene and allowed us to induce local formation of oncogenic lesions *in vivo*.

## Main

Strategies for genetic manipulation *in vivo* largely rely on site-specific DNA recombinases, with the Cre recombinase being the most widely utilized among them. To achieve conditional regulation of gene expression, Cre is often fused with an inducer-responsive-domain, facilitating recombination between loxP-flanked target DNA fragments^1,2^. In the Cre-lox mouse models, some spatiotemporal control in genome editing can be achieved by selecting a cell-type-specific promoter for Cre expression along with adjusting the chemical inducer type, dose, and timing of induction at developmental or adult stages^3–10^. However, this strategy triggers gene editing in all cells where the promoter is active, causing systemic phenotypes that often mask local effects of the gene regulation^8,11,12^. Chemical induction of engineered Cre *in vivo* also introduces additional caveats of non-specific side effects and toxicity^13–15^. Direct delivery of tissue-specific, transiently expressed Cre recombinase improves spatial and temporal regulation by restricting the induction to a specific organ but still lacks the capability to control Cre activity in a locally restricted area within the target organ^16–19^. These limitations hamper our ability to model localized biological processes *in vivo*.

Tumor initiation is a critical step of oncogenesis, but it has been challenging to investigate. In particular, development of animal models that enable spatiotemporal regulation of local tumor initiation has been impacted by the lack of suitable genome editing approaches. Primary tumors typically originate at a single site and gradually progress to stages of invasion and dissemination. However, animal models using Cre-mediated conditional oncogene expression^19–21^ or exposure to carcinogens^22^ often generate multiple tumors growing simultaneously at different locations within an organ, making it difficult to study the shifts in tumor microenvironment typically associated with clinical nascent tumors^23^. Attempting to limit the number of tumors by reducing the Cre recombinase or carcinogen dose decreases the probability of tumor formation and extends the experimental timeline, making early tumor detection time-consuming and cost prohibitive. Alternative approaches, such as direct implantation of tumor cells or tumor tissue, can generate local onset of tumors but bypass the critical early stages of tumor initiation^24^. Thus, there is a need for tumor models that can effectively simulate early human tumorigenesis while also capturing the microenvironmental crosstalk associated with tumor development.

With recent advances in optogenetic technologies, photoactivatable Cre recombinases have emerged as a promising solution for local regulation of genome editing^25,26^. An ideal optogenetic Cre for spatiotemporal control *in vivo* must exhibit a strong light-induced recombination activity, while demonstrating no background recombination in the dark. Most reported optogenetic Cre recombinases rely on light-mediated dimerization of split-Cre systems^27–33^. They often suffer from unintended spontaneous assembly and imbalanced expression levels of the two components, resulting in suboptimal performance and/or high background activity. A single-component, light-inducible Cre (LiCre) has shown very strong photoactivity in mammalian cell culture^34^. However, this system also exhibits targeted DNA binding in the dark, indicating potential background recombination activity^35^. All these limitations ultimately compromise the application of existing photoactivatable Cre recombinases for spatiotemporal regulation of genetic recombination *in vivo*.

An essential element required for *in vivo* optogenetic regulation is a system for local illumination that can offer tunable control over illumination intensity, duration, area, and timing. Tethered fiber optics^36,37^ can physically damage the targeted tissue, while the application of external illumination devices is limited due to the poor penetration of light through animal tissue, making it difficult to reach internal organs. *In vivo* illumination strategies using implantable devices overcome these limitations. The advancement of wireless µ-ILEDs controlled via near field communication (NFC) enabled reliable operation in freely moving animals^38–40^. Previous implantable optoelectronic devices have predominantly been developed for applications targeting the central nervous system^38,39,41,42^. However, their adaptability for use in other organs and peripheral nervous systems has been limited^40,43,44^. Wireless device integration with soft, moving organs such as the lungs is challenging due to the mechanical compliance and biocompatibility of the device materials. Existing devices offer limited flexibility of implantation angle, depth, relative orientation within the body, and adaptability to the animal’s motion, which can impact the performance of the device and cause permanent tissue damage. Furthermore, current approaches require high power illumination that raises concern of local tissue heating and photodamage.

Here, we report a system that overcomes the limitations of current approaches for local regulation of gene expression and tumorigenesis and demonstrates its utility in a lung tumor model. We developed a photoactivatable Cre system, E-LightR-Cre, enabling precise spatiotemporal gene editing *in vivo*. E-LightR-Cre ensures robust light-mediated regulation of gene expression with no detectable leaky activity in the dark. To achieve *in vivo* optogenetic activation, we developed wireless, battery-free, miniaturized, fully-implantable optoelectronic devices that integrate a soft, flexible, serpentine interconnect with a microscale inorganic LED (µ-ILED), mounted on a lightweight, thin, flexible, printed circuit board. A custom-designed software interface (GUI) provides real-time light control, allowing precise spatiotemporal modulation of light by controlling parameters such as duty cycle, operating duration, and power to initiate and regulate gene editing and oncogenesis locally. This wireless system ensures spatiotemporal optogenetic stimulation in animals with minimal mechanical interference, enabling reliable, continuous experiments for adjustable surgical depth and illumination area. We demonstrate the tunability of localized illumination with these devices *in vitro*, showcasing their ability to induce spatially confined gene expression and trigger oncogenic transformation in a spatially restricted population of cells. *In vivo,* intrathoracic placement of the µ-ILED over the lung enabled localized E-LightR-Cre activation, inducing local expression of a reporter gene in the illuminated area. Importantly, these implants did not affect mouse health, mobility, or lung architecture while providing stable wireless power delivery. Application of these technological advancements allowed us to initiate focal onset of lung adenocarcinoma in LSL-KRas-G12D mice. Overall, our strategy can be employed as a versatile tool for precise disease modeling and targeted gene therapy, ultimately advancing preclinical research and therapeutic innovation.

## Results

### Development of light regulated Cre recombinase demonstrating no leaky activity in the dark

Spatiotemporal control of gene editing *in vivo* requires a light-regulated Cre construct that demonstrates no background activity in the dark. We previously reported the development of light-regulated Cre recombinase LightR-Cre^45^ (Fig. 1a). To assess whether LightR-Cre possesses properties needed for application *in vivo*, we carefully evaluated regulation of LightR-Cre by blue light. We used mTmG reporter^46^ that basally expresses a membrane-targeted tdTomato and switches to expression of membrane-targeted GFP following Cre-mediated recombination (Fig. 1b). Our analysis confirmed robust activation of LightR-Cre by blue light in HEK293T cells (Fig. 1c). However, we detected a low level of background recombination in the dark when compared to the inactive mutant of LightR-Cre (Y324F ^29^, CI-LightR-Cre) (Fig. 1c, Supplementary Fig. 1a).

**Fig. 1:**
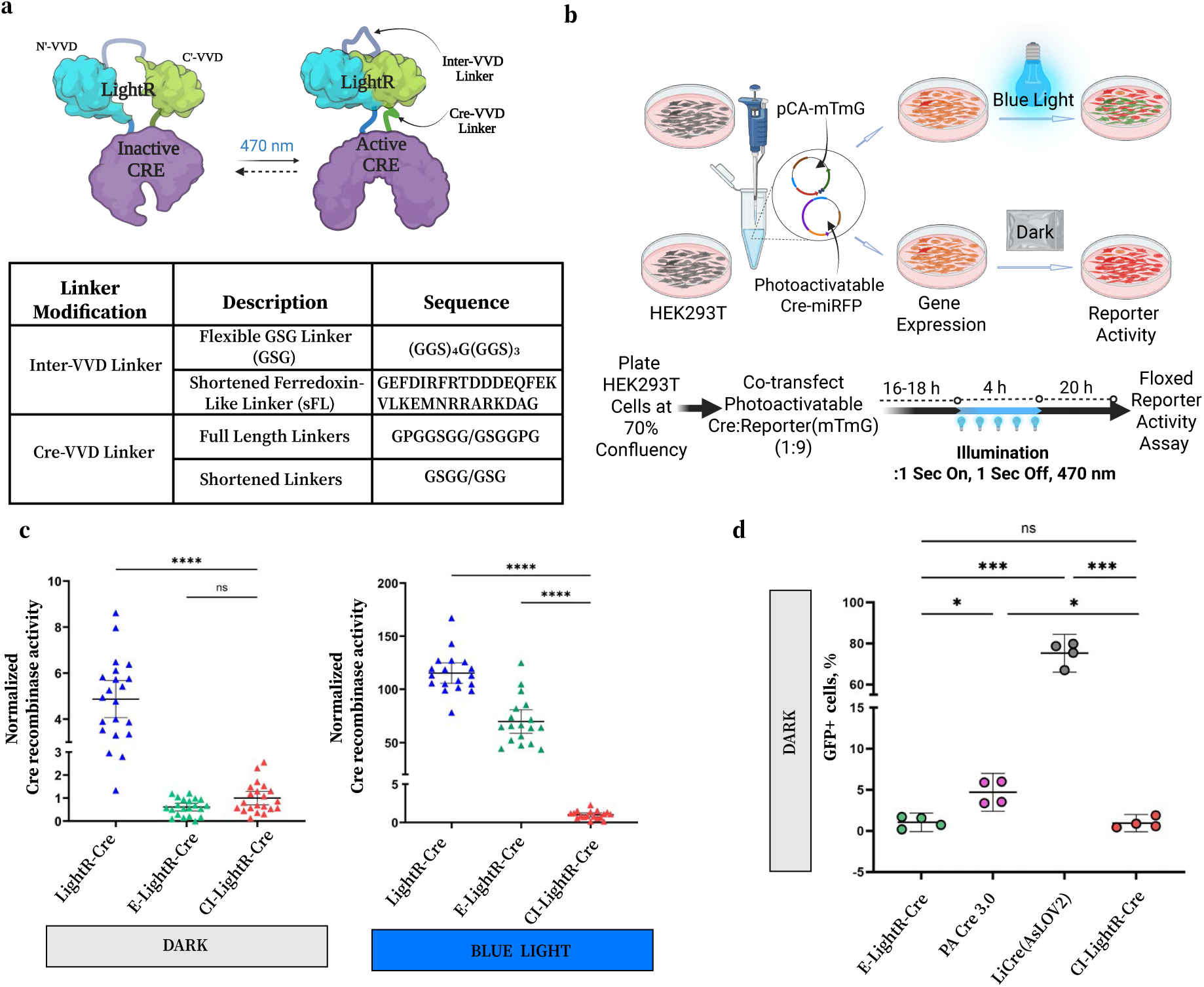
Development of light-regulated Cre recombinase demonstrating no leaky activity in the dark. **a**, Schematic representation of E-LightR-Cre design. **b**, Light-regulated Cre activity assay. HEK293T cells transiently co-transfected with miRFP670 tagged light-regulated Cre recombinase and a reporter construct (pCA mTmG) were illuminated with blue light and GFP expression was assessed either by microscopy (**c**) or flow cytometry analysis (**d**). Illustration created using BioRender (https://biorender.com). **c**, Cre recombinase activity of indicated constructs measured as the ratio of GFP-positive to tdTomato-positive cells normalized to the activity of constitutively inactive CI-LightR-Cre (Mean ± 95% confidence interval, 3 biological replicates, 7 fields of view analyzed per replicate). Statistical significance was assessed by Brown-Forsythe and Welch one-way ANOVA with Dunnett’s multiple comparisons test; ns: not significant, p > 0.05; ****: p < 0.0001; **d**, Activity of the indicated Cre constructs in the dark. Fraction of GFP expressing cells was assessed by flow cytometry analysis. The gating details are shown in Supplementary Fig. 1c. Mean ± 95% confidence interval is shown for n=4 experimental replicates. Statistical significance was assessed by Brown-Forsythe and Welch one-way ANOVA with Dunnett’s multiple comparisons test; ns: not significant, p > 0.05; *: p < 0.05; ***: p < 0.001.

Even low level of leakiness can interfere with application of light-regulated Cre *in vivo* to study chronic processes. Therefore, to eliminate the leakiness of LightR-Cre in the dark, we introduced additional modifications to the LightR domain. The LightR domain is comprised of two tandemly connected VVD photoreceptor domains that homodimerize in the blue light and dissociate in the dark, creating a light-sensitive clamp that is open in the dark and closes upon illumination (Fig. 1a). Insertion of the LightR domain into Cre at position D153 made it regulatable by light, where the open LightR clamp inactivates Cre in the dark and light-induced closing of LightR rescues Cre activity^45^. We hypothesized that the leakiness of LightR-Cre in the dark-state occurs due to spontaneous dimerization of the VVD domains, resulting from high flexibility of the connecting linkers. Thus, to develop a tightly regulated variant of LightR-Cre with no leakiness, we replaced the highly flexible 22 amino acid inter-VVD linker (GSG-linker) with a more rigid 31 amino acid sFL-linker “GEFDIRFRTDDDEQFEKVLKEMNRRARKDAG” (Fig. 1a). The sFL-linker is a shortened version of the previously reported ferredoxin-like linker peptide which demonstrates reversible unfolding and refolding upon induction and withdrawal of mechanical load, without any hysteresis^47^. We hypothesized that the increased rigidity of the sFL linker would prevent spontaneous homodimerization of the VVD domains in the dark and thus reduce unwanted leakiness of LightR-Cre. In a parallel study, we successfully utilized the sFL linker to eliminate background leakiness in a light-regulated kinase. Also, our previous results show that variation of the linkers connecting an allosteric switch domain to the target enzyme affects the efficiency of regulation^48–50^. Thus, we replaced the flexible linkers GPGGSGG and GSGGPG at the N and C termini of the LightR (Cre-VVD linkers, Fig. 1a) to shorter GSGG and GSG respectively, which can further reduce the leaky activity in the dark. We named the modified Cre as E-LightR-Cre. Fluorescent microscopy analysis of Cre-reporter activity revealed that E-LightR-Cre exhibits nearly 8-fold reduced recombination frequency in the dark, compared to the original LightR-Cre (Fig. 1c). No significant difference was observed between E-LightR-Cre and CI-LightR-Cre in the dark, indicating no detectable leakiness with E-LightR-Cre. Importantly, under illuminated conditions, E-LightR-Cre displayed strong DNA recombination activity (Fig. 1c, Supplementary Fig. 1a).

Next, the regulation of E-LightR-Cre was further compared to two other blue-light-inducible Cre recombinase systems, PA Cre 3.0^30^ and LiCre(AsLOV2)^34^, which have shown the most efficient light-mediated regulation of DNA recombination in previous reports. Flow cytometry analysis of the reporter expression in HEK293T cells revealed that the PA Cre 3.0 and the LiCre(AsLOV2) both show significant leaky activity in the dark, when compared to inactive CI-LightR-Cre (Fig. 1d and Supplementary Fig. 1b, c). E-LightR-Cre exhibited no detectable leaky activity in the dark. The light-induced activity of E-lightR-Cre was comparable to PA Cre 3.0 with no significant difference observed (Supplementary Fig. 1b). Although LiCre(ASLOV2) was found to show the highest level of light-induced activity, the pronounced leakiness in the dark renders its application *in vivo* very challenging (Fig. 1d and Supplementary Fig. 1b, c). Overall, our data demonstrate that modified E-LightR-Cre exhibits much improved regulation required for optogenetic experiments *in vivo*.

### Design and operational features of wireless, battery-free, miniaturized, fully-implantable optoelectronic devices

Previous wireless implantable optoelectronic devices designed for applications in the central nervous system have shown limited adaptability to internal organs and peripheral nervous systems, highlighting the need for technologies optimized for broader applications especially in moving soft organs such as lungs. To address these limitations, we implemented new engineering solutions to the electronic circuits, the operational scheme of the device, as well as its architecture and composition to enhance mechanical flexibility, biocompatibility, and ability to modulate the brightness and the illumination duration for localized optogenetic control of lung tissue in freely moving mice. The overall design of the implantable optoelectronic device includes an RF power harvesting system, comprised of an inductive planar coil antenna and a rectifier, coupled to a μ-ILED by a flexible trace and encapsulated in a transparent, flexible, and biocompatible materials (Fig. 2). The optoelectronic design integrates top and bottom Copper (Cu) layers surface-mounted on a 50 μm thick Polyimide (PI) substrate and encapsulated in parylene followed by a soft and flexible layer of silicone (Ecoflex; Poly(dimethylsiloxane), PDMS), adapting a planar geometry that facilitates a large-scale manufacturing process (Fig. 2a, Supplementary Fig. 2). A circular coil antenna wirelessly harvests power via magnetic inductive coupling to a resonant antenna around an animal enclosure, powered by an RF source at 13.56 MHz. The electrical components on the device include a capacitor that provides impedance matching and a Schottky diode that rectifies the received RF signals to yield a current source for the µ-ILED (Fig. 2b, Supplementary Fig. 3). A soft, flexible trace features a pair of metal lines that extend along a serpentine interconnect (24 mm L x 1.8 mm W), providing both vertical and horizontal freedom relative to the connected circular receiver coil (10 mm diameter; copper traces: 6 turns, 60 μm width, 18 μm thickness, 73 μm spacing). At the distal end, the serpentine interconnect integrates a μ-ILED (270 μm L × 220 μm W × 50 μm H), allowing for localized optogenetic stimulation. The serpentine interconnects can be fabricated in customizable lengths, supporting potential applications targeting peripheral nerves or visceral organs (Supplementary Fig. 2b). The device includes two μ-ILEDs: one emitting blue light (470 nm) at the end of the serpentine coil for optogenetic activation, and a second emitting red light (650 nm) within the receiver coil to provide a visible indicator of system activation and confirmation of device operation^51^. The miniaturized device dimensions and ultralightweight design (∼60 mg; Fig. 2b) were implemented to ensure successful implantation in adult mice without impacting their health or motion.

**Fig. 2:**
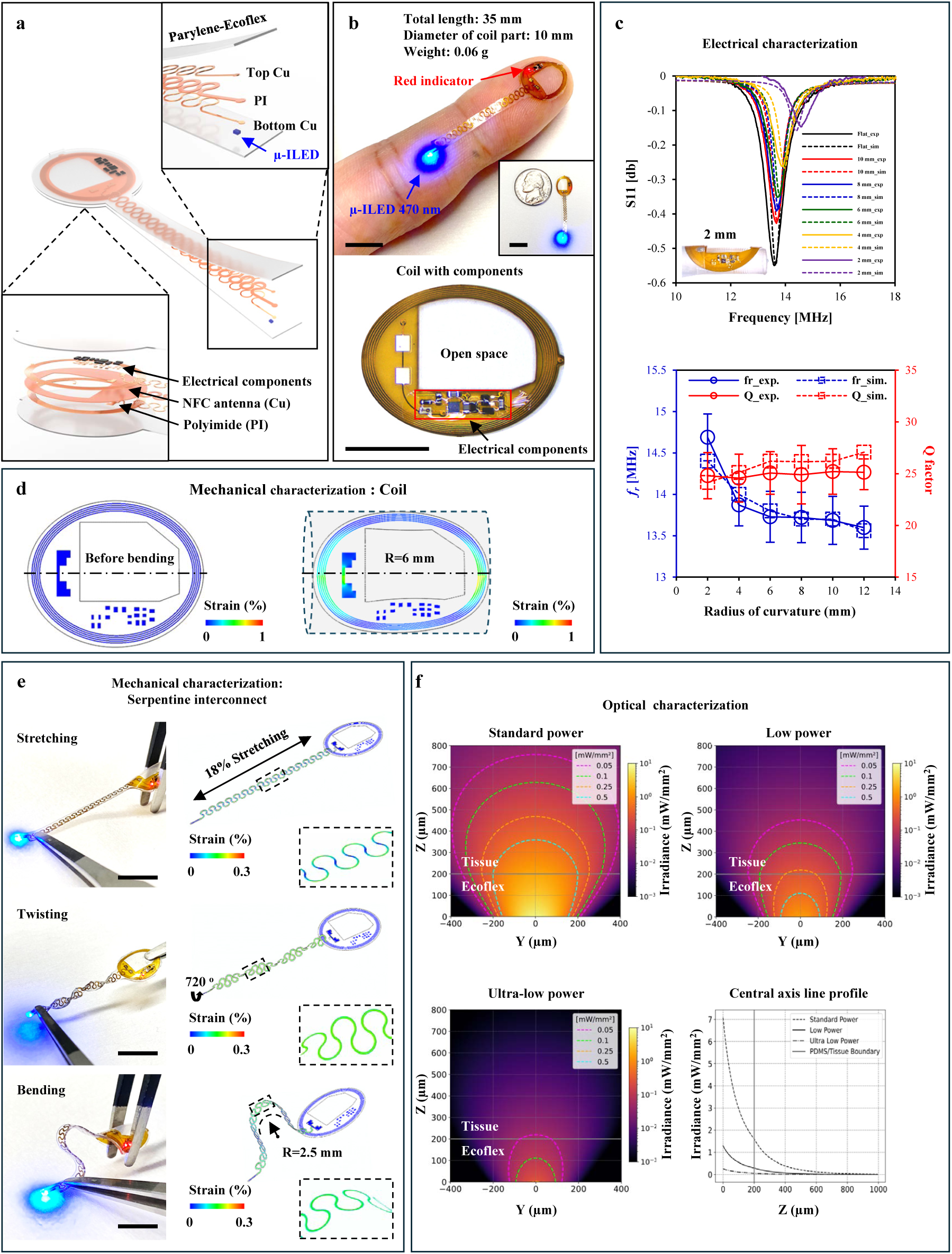
Development of wireless, battery-free, miniaturized, fully-implantable optoelectronic device and characterization of its electrical, mechanical and optical properties. **a**, Schematic illustration of the overall construction highlights a flexible serpentine interconnect with a µ-ILED at the tip end, connected to a receiver coil with matching capacitors, a rectifier, and a separate µ-ILED indicator. **b**, Top: optical image of the device showing the overall design and size, highlighting the soft, flexible, stretchable interconnect with a blue µ-ILED at the tip and a separate red indicator LED. Scale bar 1 cm. Bottom: optical image showing the top view of the coil within the electronic circuitry, including all integrated components. Scale bar 5 mm. **c**, Electrical performance analysis: Experimental (exp) and simulated (sim) antenna performance of the coil as a function of bending curvature. Top: Simulated reflection scattering parameter (S11) versus antenna frequency at different curvature radii. Bottom: Comparison of the experimental and simulated values of the resonant frequency (*f_r_*) and Q factor of the antenna at different radii of curvature. **d**, **e**; Mechanical performance analysis: **d**, Device coil: Distribution of effective strain in the copper coil of the device body, before (Left) and after (Right) bending, with a radius of curvature of R=6 mm. **e**, Serpentine interconnect: Images of the activated devices and corresponding Finite Element Analysis results in stretching, twisting, and bending mode. Scale bar 1 cm. The color map represents effective strain distribution. **f**, Irradiance profiles of blue μ-ILEDs with varying current-limiting resistors for spatiotemporally confined illumination. Each condition is accompanied by a central axis irradiance line plot.

### Electrical and mechanical properties of the optoelectronic devices

Reliable operation of the device under mechanical stress exerted by freely moving animals is critical for successful applications in mouse models. Thus, we evaluated the key electrical and mechanical properties of the devices.

The electrical characterization of the device coil reveals that experimental measurements of scattering parameters (S11), resonant frequency (*f_r_)*, and Q factor at different radii of curvature match the simulated performance of the antenna. These results demonstrate that the device maintains reliable functionality and consistent optical stimulation even under significant mechanical deformation, both in experiments and simulations (up to a radius of curvature of 4 mm), thereby ensuring stable performance during the natural locomotion of small animals (Fig. 2c).

To evaluate the mechanical performance of the device coil, we quantified strain distribution in the metal traces of the coil antenna when bent to a 6 mm radius. Our simulation results show that the metal traces experience strain levels exceeding the yield point (0.3%) but remaining well below the fracture limit for copper (<1%) (Fig. 2d). We also assessed the mechanical performance of the serpentine interconnects encapsulated by a soft silicone layer under various deformation conditions. Finite element analysis (FEA) was used to simulate stretching (up to 18%), bending (to a radius of curvature R = 2.5 mm), and twisting (up to 720 °) of the serpentine interconnect. This analysis demonstrates that the serpentine coil could accommodate extensive mechanical deformations without yielding, as the strain in the metal remains below 0.3% (Fig. 2e). Thus, the mechanical compliance aligns with the types of deformation experienced by soft biological tissues^40,52^. Additional mechanical tests, including tensile and cyclic testing, demonstrate that the encapsulating layers provide effective barrier properties, ensuring sufficient operational stability and stretchability for implantation *in vivo*. The flexible serpentine interconnects accommodate approximately 210% strain without causing mechanical or electrical failure of the embedded conductive Cu traces (Supplementary Fig. 4a, b). They also maintain structural integrity and functionality under a wide range of mechanical deformations caused by animal locomotion, enduring from −40% bending to +50% stretching over 10,000 repetitive cycles without degradation in performance (Supplementary Fig. 4c). Taken together, these results demonstrate that the device offers reliable functionality under mechanical deformation, allowing for bending, motion and confirmational flexibility without compromising its ability to harvest wireless energy and allowing for long-term animal experiments with minimal interference.

### Optical properties of the optoelectronic devices

The optoelectronic devices were modified to operate the blue μ-ILEDs at varying light intensities to spatiotemporally confine the illumination. To achieve this, we utilized three resistors (Rs) with different current-limiting capacity (Rs=200 Ω : 580 µA, 2 kΩ :135 µA, 10 kΩ : 37 µA), and named the devices as Standard_power (*I_o_*=165 µW; previously used in optoelectronic devices and standard for blue μ-ILEDs), Low_power (*I_o_*=30 µW), and Ultra-low_power (*I_o_*=6 µW). Simulation of irradiance profiles in mouse lungs shows that the predicted penetrance of the tissue and illuminated volume of low and ultra-low power devices is dramatically reduced and thus should enable more localized optogenetic regulation (Fig. 2f, Table 1) and mitigate or even eliminate any potential for thermal or photodamage.

### Local regulation of E-LightR-Cre using the optoelectronic device *in vitro*

One of the advantages of optogenetic tools is the capability to vary the zone of activation depending on the specific goals of the experiment. Thus, we evaluated whether we could control the area of E-LightR-Cre activation by modulating illumination intensity of µ-ILED devices and duration of illumination. We employed the Low_power (*I_o_*=30 µW) and Ultra-low_power (*I_o_*=6 µW) µ-ILED devices, to evaluate the area of E-lightR-Cre activation in HEK293T cells co-transfected with Floxed-STOP-mCherry reporter construct^29^ (Fig. 3a). Our data demonstrate that by adjusting illumination intensity and duration, we can control the area of E-LightR-Cre activation (Fig. 3b, c) with a minimum average activation radius of less than 2 mm from the center of illumination (Fig. 3c).

**Fig. 3:**
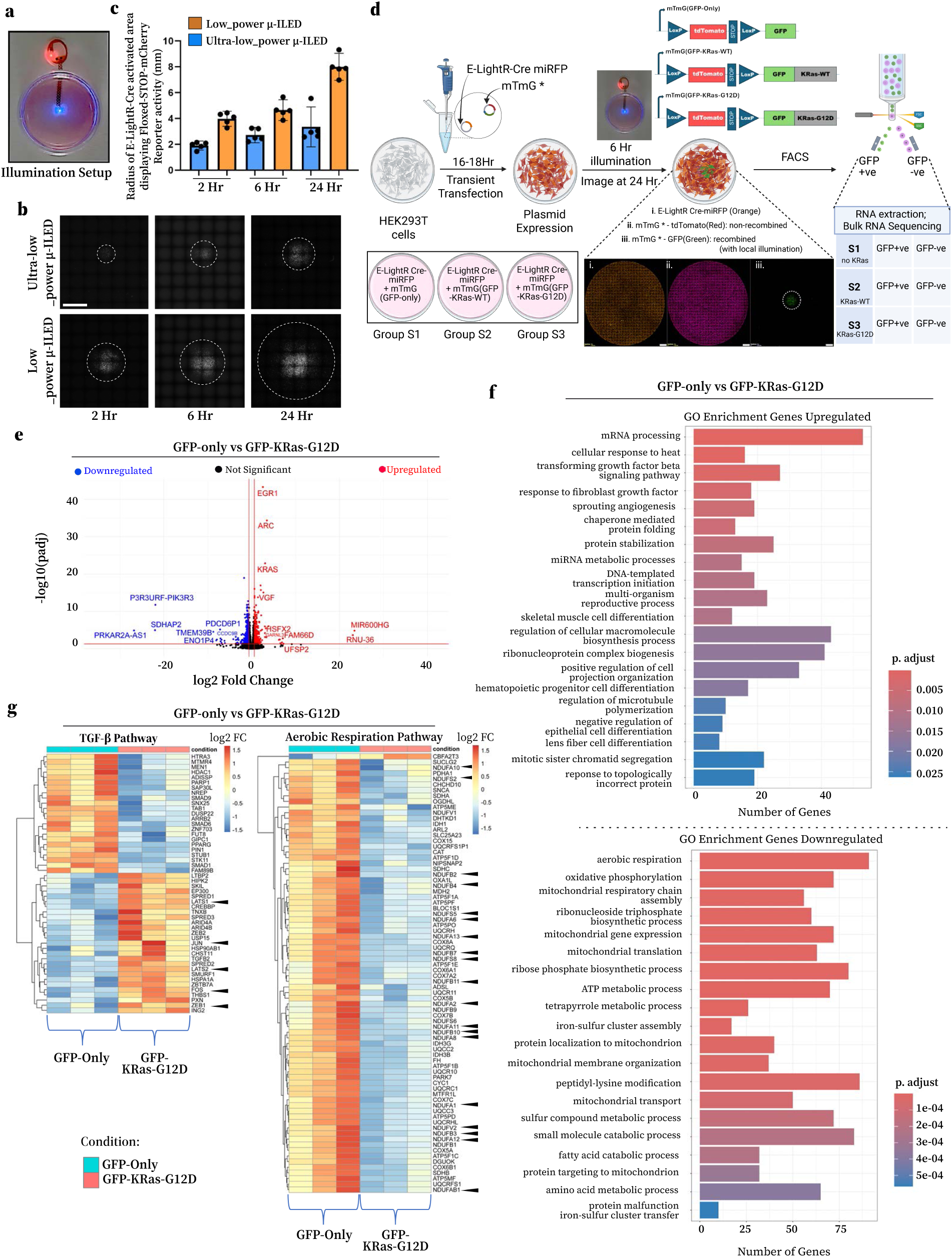
Characterization of E-LightR-Cre mediated spatiotemporal control of genome editing using a wireless µ-ILED *in vitro*. **a**, *In vitro* setup for localized illumination, using wireless µ-ILED. The devices were attached beneath a 35mm culture dish with the illuminating tip positioned at the center. **b**, **c**; Effect of different illumination intensity and duration on area of activation. HEK293T cells co-transfected with E-LightR-Cre-miRFP670 and a Floxed-STOP-mCherry reporter were illuminated with specified µ-ILEDs for defined durations, then incubated in the dark to complete 24 hours post-illumination. **b**, Epifluorescence wide-field images showing area of mCherry expression (outlined with dotted circles) under the defined illumination conditions. Scale bar 5 mm. **c**, Average radius of E-LightR-Cre induced mCherry expression zone. Data represented as mean ± 95% confidence interval, n=5. **d**, Experimental design for evaluating spatiotemporal regulation of KRas-G12D expression *in vitro*. HEK293T cells co-transfected with E-LightR-Cre-miRFP670 and one of the three indicated reporter constructs were locally illuminated using Low_power µ-ILED device (470 nm; 10 W, 10 Hz, 20% duty cycle) for 6 hours. Following illumination, cells were incubated in the dark for 18 hours. GFP-positive and GFP-negative cells from each group were sorted by flow cytometry and used for total RNA extraction and bulk RNA sequencing analysis (n=3 in each group). Illustration created using BioRender (https://biorender.com). **e**, Volcano plot of the top differentially expressed genes comparing GFP-only with GFP-KRas-G12D expressing cells. Top 17 differentially expressed genes are annotated. **f**, Gene Ontology (GO) enrichment analysis of differentially upregulated (Top) or downregulated (Bottom) pathways comparing between GFP-only and GFP-KRas-G12D expressing cells. Significance criteria: adj. p-value < 0.05. **g**, Heatmaps of differentially expressed genes under TGF-β Pathway (Left) and Aerobic Respiration Pathway (Right). Significance criteria: adj. p-value < 0.05. Color-coded scale corresponds to indicated fold change in gene expression (log2 FC).

Next, we assessed the ability of E-LightR-Cre to induce specific signaling and oncogenic transformation in a locally illuminated population of cells. We generated reporter constructs that express membrane targeted tdTomato flanked by loxP sites and followed by GFP-tagged wild-type KRas or G12D mutant (Fig. 3d). GFP-tagged wild type KRas or its oncogenic mutant (G12D) will be expressed only in cells where E-LightR-Cre is activated. Using the optoelectronic device µ-ILEDs, we induced expression of GFP-KRas-G12D or GFP-KRas-WT in a local population of HEK293T cells lacking intrinsic KRas mutations. Cells transfected with unmodified mTmG reporter construct served as a negative control (Fig. 3d). After 6 hours of local illumination followed by 18 hours of incubation in the dark, GFP-positive cells were observed in the illuminated regions, while GFP-negative/tdTomato-positive cells remained at the non-illuminated periphery. To evaluate the effect of oncogenic KRas, GFP-positive and GFP-negative cells from each group were sorted using fluorescence-activated cell sorting (FACS), and total RNA was extracted from the sorted cells for bulk RNA sequencing analysis (Fig. 3d). Principle component analysis demonstrated clustering of all GFP-negative cells and visualized individual clusters of GFP-only, GFP-KRas-WT and GFP-KRas-G12D expressing cells (Supplementary Fig. 5a). We identified 3,798 and 2,106 early responding genes differentially expressed in the GFP-KRas-G12D cells compared to GFP-only and GFP-KRas-WT cells, respectively (adj. p-value < 0.05; Table 2). This included an upregulation of 1,219 genes and downregulation of 2,579 genes relative to the GFP-only cells, and the upregulation of 941 genes and downregulation of 1,165 genes relative to the GFP-KRas-WT cells (Table 2).

Among the top differentially expressed genes in the GFP-KRas-G12D positive cells were *egr1*, *arc* and *vgf* (Fig. 3e, Supplementary Fig. 5b). These genes have been previously implicated as downstream targets of oncogenic Ras signaling^53–55^.. Gene ontology (GO) enrichment analysis of the GFP positive population in the GFP-KRas-G12D cells compared to GFP-only cells revealed upregulation of TGF-β and FGF pathways along with downregulation of aerobic respiration and oxidative phosphorylation pathways (Fig. 3f), consistent with previous findings in KRas transformed cells^56–58^. The differentially expressed genes within the identified biological pathways are shown in representative heatmap diagrams (Fig. 3g). Analysis of TGF-β pathway genes revealed upregulation of *jun* and *fos,* consistent with activation of early oncogenic signaling induced by KRas-G12D^59^. We also observed upregulation of *zeb1*, *lats1* and *lats2* genes, in line with previous reports identifying them as downstream regulators in Ras-driven cancers^60–63^. Downregulation of the aerobic respiration pathway, marked by significant reductions in the mitochondrial complex I-IV genes including a major decrease in major complex-I subunit genes (*ndufa*, *ndufb*, *ndufs*, *ndufv*, and *ndufab1*), suggests a metabolic switch within 24 hours of oncogenic KRas-G12D expression (Fig. 3g). This aligns with previous reports of suppressed mitochondrial complex-I activity and mitochondrial dysfunction induced by mutant KRas^58,64^. When compared to the GFP-KRas-WT, the GFP-KRas-G12D-expressing cells display a similar trend in upregulation of TGF-β signaling along with downregulation of aerobic respiration and mitochondrial complex genes, thereby promoting early oncogenic transformation and metabolic reprogramming (Supplementary Fig. 5c, d). Together, these findings demonstrate that E-LightR-Cre induces KRas-G12D driven tumorigenic signaling within 24 hours of locally illuminating cells.

We also analyzed the effect of the KRas-G12D-transformed cells on their non-transformed GFP-negative neighbors in the same samples. We observed significant differential expression of several genes in the non-illuminated cells that did not undergo recombination. In the GFP-negative population of the mTmG(GFP-KRas-G12D) group, 13 genes were differentially expressed when compared to the mTmG(GFP-only) cells (adj. p-value < 0.05; Table 2) and 5 genes were differentially expressed when compared to the mTmG(GFP-KRas-WT) cells (adj. p-value < 0.05; Table 3). These data suggest that KRas-G12D expression rapidly (within 24 hours) reprograms cells, and importantly, also reprograms the non-transformed neighbors.

### Device implantation and biocompatibility

To achieve optogenetic regulation of E-LightR-Cre in mouse lungs, we implanted the wireless µ-ILED device by posterolateral thoracotomy to place the illuminating tip over the left lung lobe. The circular receiver coil for wireless NFC signal was inserted subcutaneously along the back of the mouse, while the stretchable serpentine metallic trace, coated with implantation-compatible Ecoflex/PDMS, was inserted into the thoracic cavity between the 6^th^ and the 7^th^ ribs, positioning the µ-ILED over the left lobe of the lung (See Methods, Fig. 4a, Supplementary Fig. 6a). Successful implantation with optimal positioning of the device was confirmed by respiratory gated live and post-mortem high resolution, whole body micro-CT imaging, 4 weeks post-surgery (Fig. 4b). The device was retained at its desired position in all mice (Supplementary Fig. 6b) at their respective experimental end points (up to 16 weeks post-surgery). Assessment of device biocompatibility indicated no signs of lung damage at the device contact site (Fig. 4c). The body weight of the animals showed no statistically significant differences in either males (n=3) or females (n=3) when comparing preoperative day 1 to postoperative time points at 3 days, 1 week, and 2 weeks (Fig. 4d). No major postoperative complications were observed in the implanted animals. We also analyzed the impact of the implants on locomotor behavior of the mice comparing their activity levels. The results showed no significant differences in average traveled distance or average velocity at 1 week or 4 weeks post-surgery compared to the day before the surgical procedure (Fig. 4e, f). Collectively, these results suggest that the implanted devices do not cause detectable damage to the mouse lungs and do not interfere with natural movement or normal activity levels of mice.

**Fig. 4:**
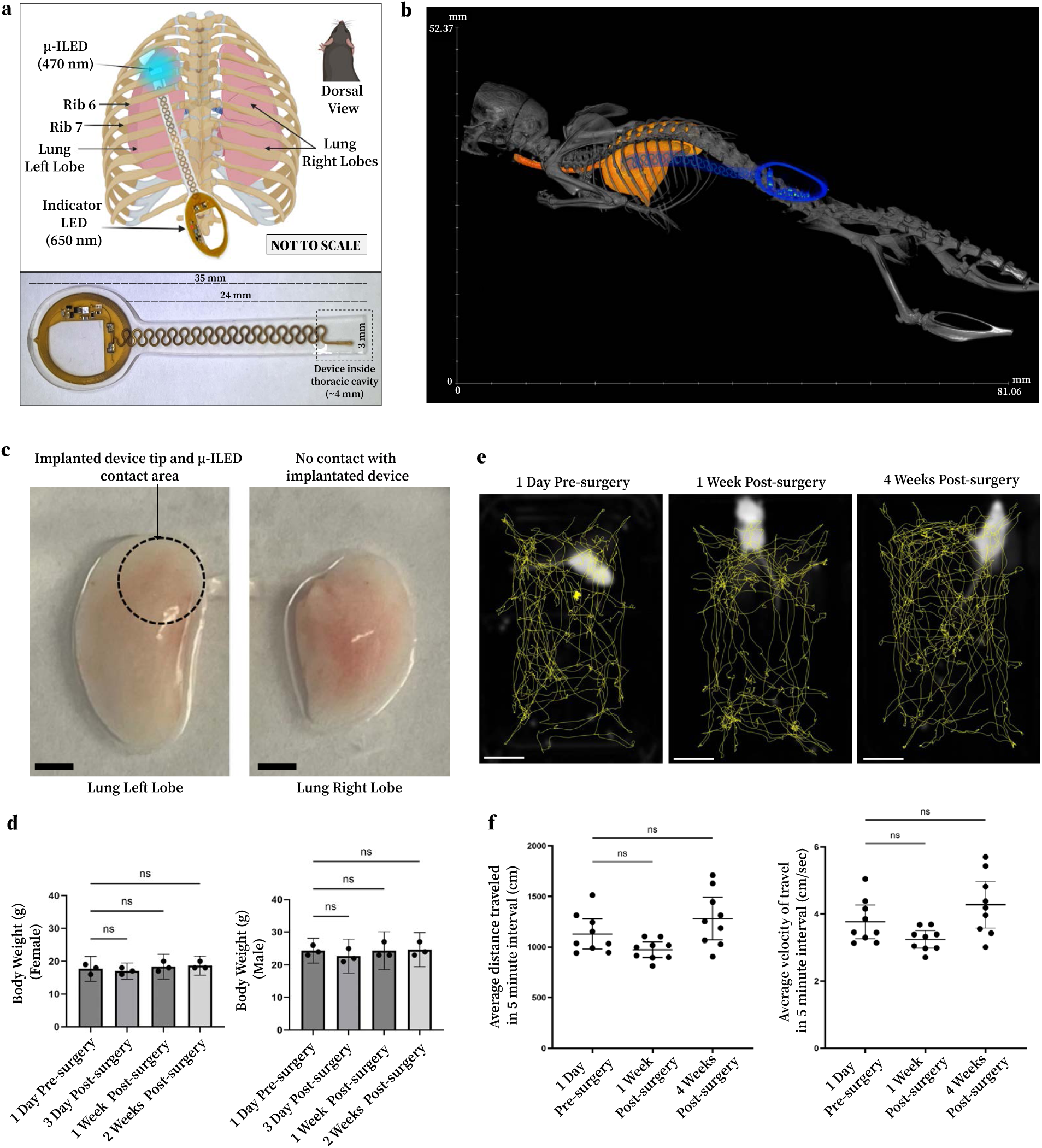
Surgical implantation of optoelectronic µ-ILED device. **a**, Illustration of posterolateral, back-mounted, intrathoracic implantation of peripheral wireless μ-ILED device in a mouse. The device tip is inserted into thoracic cavity to position the µ-ILED (470 nm) over the left lobe of the mouse lung. The inserted tip (∼4 mm) is outlined with a dotted line. Illustration created in BioRender (https://biorender.com) with additional modifications. **b**, Respiratory-gated *in vivo* micro-CT imaging showing a back-mounted intrathoracic implantation of a wireless μ-ILED device. The stretchable serpentine trace is inserted into the thoracic cavity between 6th and 7th ribs positioning the illuminating tip over the left lobe of the lung. **c**, Representative photograph of mouse left lung lobe and one of the four right lung lobes. The dotted circle indicates the area near the implanted μ-ILED tip, showing intact lung surface. Scale bars 2 mm. **d**, Mouse body weights at the indicated time before and after device implantation surgery. Statistical analysis was conducted using paired, repeated measures one-way ANOVA followed by Tukey’s multiple comparisons test. Data are presented as mean ± 95% confidence interval (n=3 males, 3 females, ns: non-significant, p > 0.49. **e**, Representative movement tracks of individual mice recorded over a five-minute period in an open-field environment. Thresholded mouse images shown as white contours mark the starting points of motion tracking. Scale bars 5 mm. **f**, Average distance traveled (Left) and Average velocity (Right) measured over a five-minute period in a rectangular open-field test at the indicated time points before and after surgery. Statistical significance was assessed by Brown-Forsythe and Welch one-way ANOVA with Dunnett’s multiple comparisons test; ns: not significant, p > 0.05. Data are presented as mean ± 95% confidence interval: n=3 five-minute time intervals for each time point, per mouse.

### Local control of gene expression in mouse lungs using E-LightR-Cre and wireless µ-ILED

To assess the spatial control of DNA recombination *in vivo*, we tested local regulation of E-LightR-Cre in the lungs of ROSA26-loxP-STOP-loxP(LSL)-tdTomato reporter mice^65^. Following implantation of the Low_power µ-ILED device, adenovirus expressing either E-LightR-Cre or unmodified WT-Cre was delivered endotracheally for whole-lung distribution. After 24 hours, mice were placed within a wireless RF chamber and the left lung was locally illuminated by the implanted µ-ILED (See Methods, Supplementary Fig. 6c). Following 10 days post-illumination, mouse lungs were harvested, cleared using the ‘Fast-3D clear’ protocol^66^ adapted for lung tissue, and analyzed using light sheet microscopy^67^ (Fig. 5a_i). Spatially restricted activation of E-LightR-Cre induced localized tdTomato expression within the confined area of the left lobe (Fig. 5b, Supplementary Fig. 7a i,ii). In contrast, the unmodified WT-Cre induced recombination across the entire left lobe of the lung (Supplementary Fig. 7b), further demonstrating the efficiency of the adenovirus delivery and its distribution in the mouse lungs. Overall, our data show that we achieved tightly regulated local control of gene expression *in vivo*.

**Fig. 5:**
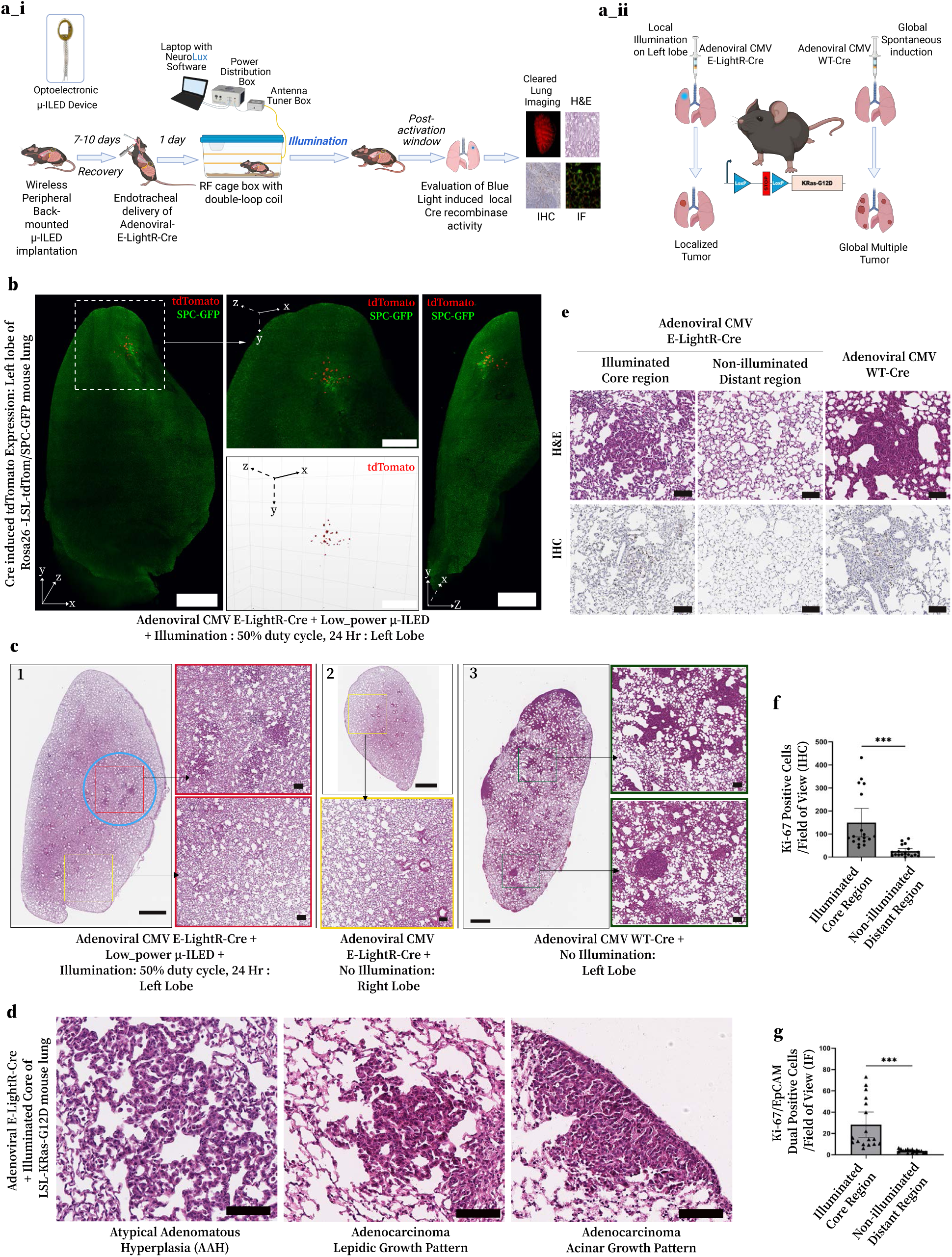
Light-mediated local genome editing and spatially restricted tumor initiation in mouse lungs. **a**, **a_i**, Schematic *in vivo* workflow for evaluating spatiotemporal gene regulation using E-LightR-Cre and wireless μ-ILEDs. **a_ii**, Schematic study design comparing global versus local light-controlled tumor stimulation. **b**, Representative LSFM 3D micrographs of cleared left lung from Rosa26-LSL tdTomato/SPC-GFP mouse, 10 days after Low_power-μ-ILED illumination following adenoviral E-LightR-Cre delivery, showing tdTomato expression (red) over constitutive SPC-GFP (green). Local activation radius (95% signal area) ≤ 0.5 mm. Scale bars: 1 mm for whole-lobe images, 500 µm for insets. **c**, Representative H&E Images of LSL-KRas-G12D mouse lung sections 8 weeks after adenoviral Cre induction. Panel 1, 2; representative illuminated left lung (1) and non-illuminated right lung (2) sections from mouse that received adenoviral E-LightR-Cre and local illumination. Illuminated area on the left lung outlined by a blue circle. Zoomed-in images show illuminated (red rectangle) and non-illuminated distant (yellow rectangle) regions. Panel 3; representative left lung section from mouse that received unmodified adenoviral WT-Cre. Scale bars; 1 mm for whole-lobe sections, 100 µm for zoomed-in regions. **d**, Representative H&E images of different types of local lesions obtained 8-14 weeks post μ-ILED-induced E-LightR-Cre activation. **e**, Representative lung sections from the samples in (**c**), with H&E staining and Ki-67 IHC staining from consecutive sections. Scale bars 100 μm. **f**, Quantification of Ki-67-positive cells per field of view. Data presented as mean ± 95% confidence interval. N=18 fields of view across three mice. **g**, Quantification of Ki67-EpCAM dual-positive cells in each field of view from indicated lung regions, co-stained for Ki-67 and EpCAM. Data presented as mean ± 95% confidence interval. N=17 fields of view across three mice. Statistical significance (**f**, **g**) was determined by unpaired two-tailed t-test with Welch’s correction (***, p < 0.001).

### Local stimulation of tumor initiation in mouse lungs

Local regulation of tumor initiation in mice remains a significant challenge, especially in lung cancer models. Our optogenetic approach offers a promising solution to overcome these limitations. To evaluate this, we utilized the widely used LSL-KRas-G12D mouse model, a common tool for studying lung cancer development^19^. We tested whether optogenetic control over the expression of the oncogenic KRas-G12D mutant could initiate the development of lung tumors in a local area of the lungs (Fig. 5a_ii). Following implantation of the Low_power µ-ILED device and endotracheal delivery of adenoviral E-LightR-Cre, the left lobe of the lung was illuminated locally, and the formation of tumors was assessed by histological analysis. Eight weeks after induction, we observed localized neoplasia in the illuminated area of the left lobe, histopathologically characterized as foci of atypical adenomatous hyperplasia to lepidic adenocarcinoma (Fig. 5c, d; Supplementary Fig. 8a). Importantly, we found no signs of neoplasia in the distant areas on the left lobe or on the right lung lobes (Fig. 5c, Supplementary Fig. 8a). Illumination with an Ultra-low_power µ-ILED device stimulated formation of a single local lesion of acinar type advanced adenocarcinoma which appeared at fourteen weeks after induction (Fig. 5d, Supplementary Fig. 8b). These findings suggest that E-LightR-Cre induced local tumor initiation can be modulated by light intensity. Immunohistochemistry analysis revealed a significantly higher number of Ki-67-positive proliferating cells within the light-activated neoplastic mass, compared to the surrounding normal lung tissue, indicating increased proliferation within locally induced tumors (Fig. 5e, f). Immunofluorescence staining further demonstrated elevated EpCAM expression in histopathologically identified neoplastic regions relative to adjacent normal tissue, confirming the epithelial origin of the tumors^68^. Additionally, the increased presence of dual-positive Ki-67/EpCAM-high cells in the light-activated core, compared to distant lung areas, provides further support for the epithelial origin and proliferative nature of the light-induced neoplastic lesions (Fig. 5g, Supplementary Fig. 8c). In contrast, endotracheal delivery of adenovirus expressing unmodified WT-Cre led to the formation of multiple adenomatous tumors at different stages of growth throughout the lungs within 8 weeks (Fig. 5c, e; Supplementary Fig. 8c). These data suggest that locally induced lung tumors more accurately recapitulate the progression of lung tumors initiated from a single location, providing an improved model to study early events of lung cancer.

## Discussion

Despite the promises of optogenetics in spatiotemporal control of genetic modifications, its application in animal models has been hampered by current methodological limitations. Our study overcomes these by developing an optimized E-LightR-Cre recombinase and integrating it with an implantable wireless optoelectronics system for precise spatiotemporal gene editing *in vivo*. We demonstrated that E-LightR-Cre possesses the characteristics critical for *in vivo* applications: 1) no detectable leaky activity in the dark and 2) robust activation upon illumination with blue light, thereby preventing unwanted random recombination events that could complicate the analysis. Development of advanced optoelectronic devices and a precise implantation procedure allowed localized E-LightR-Cre activation *in vivo*, without causing tissue damage. With this, we demonstrated local recombination through reporter gene expression and localized initiation of tumor formation, within the mouse lungs. Our technology thus enables light-guided *in vivo* regulation of genome editing in both time and space, opening new possibilities for targeted genetic interventions.

Comparison to other optogenetic Cre systems shows that E-LightR-Cre has significantly lower background activity (Fig. 1d, Supplementary Fig. 1b, c). Two-component systems like Magnet-based PA Cre 3.0 often suffer from imbalanced expression or incomplete cleavage, leading to non-specific background activity^28,30^. While LiCre(AsLOV2) overcomes the limitations of split systems and demonstrates strong photoactivity, it exhibits targeted DNA binding in the dark, as previously reported^35^, and shows a high dark-state leakiness in our results (Fig. 1d). This makes it less suitable for extended duration of *in vivo* experiments, where unintended recombination can accumulate over time. Thus, E-LightR-Cre represents a more reliable alternative for light-regulated Cre-recombination *in vivo*.

One of the challenges of blue-light guided *in vivo* optogenetics, has been the limited depth of light penetration using externally applied illumination sources, which is especially problematic when targeting internal organs, particularly within the intrathoracic chamber. Due to constant movement and delicate tissue architecture, the lungs remain among the most challenging organs for achieving precise local optogenetic regulation. To overcome this limitation, we introduced wireless fully-implantable optoelectronic devices with a re-engineered electronic circuitry, and mechanical, optical and operational properties, that are capable of localized illumination of mouse lungs in the intrathoracic compartment, without compromising animal mobility, weight, general health, survival or tissue integrity at the point of contact (Fig. 2, 4). The device stays active in the mouse and outlives the experimental window. Wireless, implantable μ-ILED devices have been explored for preclinical and clinical studies over the past decade^38–44,52,69,70^. However, to the best of our knowledge, this is the first report demonstrating optogenetic manipulation in the lungs. Successful regulation in such a delicate organ suggests that this approach can be broadly adaptable for a wide range of *in vivo* applications, including those with less challenging requirements.

With regulated illumination fields (Fig. 2f), the μ-ILEDs with reduced powers enabled local E-LightR-Cre activation to achieve a fine precision in spatiotemporally restricted Cre-reporter activity both *in vitro* (Fig. 3 a-c) and *in vivo* (Fig. 5b, Supplementary Fig. 7a). The ability to generate such spatial heterogeneity through localized gene manipulation offers a powerful platform for studying complex cellular environments. Harnessing the specificity of E-LightR-Cre, we selectively restricted the KRas-G12D driver mutation to a defined local group of cells within an otherwise normal cell population. Light-guided, locally restricted expression of oncogenic KRas-G12D in a subpopulation of HEK293T cells *in vitro* revealed significant rewiring of gene expression pathways within 24 hours, indicating broad and rapid reprograming of the transcriptome. We identified upregulation of TGF-β pathway and downregulation of aerobic respiration and oxidative phosphorylation (OXPHOS) pathways in the local GFP-KRas-G12D expressing cells compared to GFP-KRas-WT or GFP-only cells (Fig. 3d-g, Supplementary Fig. 5), consistent with the established KRas-driven transformation events^56,58^.

Within TGF-β pathway, we observed increased expression of both oncogenic and tumor suppressor gene transcripts. While the oncogenes signify tumor-initiating signaling, the increased expression of tumor suppressors within 24 hours of KRas-G12D induction aligns well with the premalignant role of TGF-β signaling in early tumor development before transitioning to support epithelial-mesenchymal transition and metastasis^54,71–76^. Similarly, OXPHOS dependency also varies widely across cancer types, stages, and microenvironments^77–81^ . Our results showing downregulation of OXPHOS pathway upon KRas-G12D induction may indicate a metabolic shift associated with early premalignant signaling synergizing with TGF-β tumor suppressor effects. Alongside spatially restricted signaling in the illuminated area, the small-scale differential gene expression in the non-recombined periphery of the KRas-G12D transformed cells likely represents the early responses of the tumor environment to the transformed cells (Table 2, 3). Overall, optogenetic regulation of locally restricted KRas-G12D signaling offers a valuable tool to investigate how localized somatic mutations prime the environment through cell-fate determining signaling and metabolic shifts ultimately driving tumorigenesis^82^.

Consistent with previous findings, the unmodified WT-Cre recombinase induced widespread development of advanced lung tumors^11,19^ within 8 weeks (Fig. 5c). Such rapidly progressing multi-tumor models may not fully recapitulate the early stages of disease development. In contrast, our novel approach involved the application of the µ-ILED devices to activate E-LightR-Cre. This induced localized oncogenic KRas-G12D expression in mouse lungs and led to the development of fewer and slower-progressing focal lesions. These lesions predominantly consisted of atypical adenomatous hyperplasia, with fewer instances of lepidic and acinar adenocarcinoma observed at 8–14 weeks (Fig 5c, d; Supplementary Fig. 8a, b). This suggests that when KRas-G12D was induced in a small subset of cells, tumor initiation was notably impaired, likely due to a stronger anti-tumorigenic response from the surrounding tissue. This is consistent with a recent observation that limiting KRas-G12D expression in the mouse pancreas allows surrounding healthy cells to outcompete and eliminate transformed cells from the tissue^83^. Therefore, local mutations must overcome greater competition from the surrounding environment to reach the tipping point and generate tumors. By enabling precise spatial restriction of KRas-G12D expression, our model provides a useful model for studying the early cellular and microenvironmental processes that influence lung tumor formation.

In conclusion, this study introduces E-LightR-Cre as a transformative tool for genome editing, demonstrating no detectable leakiness, precise activation, and *in vivo* compatibility. The integration of flexible wireless µ-ILED systems enables spatiotemporally regulated optogenetic manipulation in the lungs and potentially other internal organs, that are inaccessible by external illumination or peripheral implantation. Along with providing a novel model system for studying spatiotemporally restricted tumor initiation and progression, this approach establishes a versatile platform for investigating complex biological systems in a precisely controlled spatial context.

## Methods

### Molecular cloning of DNA constructs

Original mammalian codon-optimized gene, encoding the LightR-Cre^45^ was amplified from our previously published construct (Addgene plasmid # 162158 ; http://n2t.net/addgene:162158 ; RRID:Addgene_162158). To obtain E-LightR-Cre, the inter-VVD linker and the linkers between Cre and LightR domain (Cre-VVD linker) were replaced using modified site-directed mutagenesis, as previously described^49^. E-LightR-Cre-miRFP670 construct was generated in the pN1 vector backbone, same as the original LightR-Cre^45^. Amino acid sequence alterations involved in linker modifications are described in Fig. 1a. To generate a constitutively inactive CI-LightR-Cre construct, a single point mutation (Y324F ^29^ – in WT Cre) was introduced into the original LightR-Cre via site-directed mutagenesis.

PA Cre 3.0^30^ in lentiviral backbone (Lv-SD-PA-Cre-nMag_Opti) was obtained as a gift from Dr. Masayuki Yazawa and LiCre(AslOv2)^34^ (Alternative names: AsLOV2-CreE340AD341A; pGY577) was a gift from Dr. Gaël Yvert (Addgene plasmid # 166663; http://n2t.net/addgene:166663; RRID:Addgene_166663). Both constructs were subcloned into pN1vector, in-frame with a C terminal miRFP670 tag, to match the backbone and the promoter strength to E-LightR-Cre-miRFP670 construct. The cloning procedure involved generation of a megaprimer followed by modified site directed mutagenesis as previously described^49^. The reporter plasmid pCA-mTmG was a gift from Dr. Liqun Luo (Addgene plasmid # 26123; http://n2t.net/addgene:26123; RRID:Addgene_26123), and the reporter plasmid pcDNA3.1_Floxed-STOP-mCherry was a gift from Dr. Moritoshi Sato (Addgene plasmid # 122963; http://n2t.net/addgene:122963; RRID:Addgene_122963). The pCA-mTmG(GFP-Kras-WT) and pCA-mTmG(GFP-Kras-G12D) reporter plasmids were cloned following the protocol for generation of megaprimers and subsequent modified site directed mutagenesis^49^. Briefly, the KRas-WT and KRas-G12D genes were amplified as megaprimers from pBabe-KRas-WT (Addgene plasmid # 75282 ; http://n2t.net/addgene:75282 ; RRID:Addgene_75282) and pBabe-KRas G12D (Addgene plasmid # 58902 ; http://n2t.net/addgene:58902 ; RRID:Addgene_58902), obtained as gifts from Dr. Channing Der. Using a modified site-directed mutagenesis strategy, the megaprimers were subsequently inserted into the pCA-mTmG vector, C terminal to GFP, to generate GFP-Kras-WT or GFP-Kras-G12D fusion genes^49^.

E-LightR-Cre-miRFP670 was further cloned into pShuttle plasmid^84^. For cloning, E-LightR-Cre-miRFP pN1 plasmid was used as a template to PCR-amplify E-LightR-Cre-miRFP gene, with primers that introduced restriction sites NotI and HindIII on the 5’ and 3’ end respectively. This PCR product and the pShuttle plasmid were digested with NotI and HindIII and then ligated to generate E-LightR-Cre-miRFP-pShuttle construct. Adenovirus production and amplification from the E-LightR-Cre-miRFP-pShuttle construct were done at the CCVR-RRC Viral Vector Core at the University of Illinois Chicago (UIC) to obtain adenovirus expressing E-LightR-Cre-miRFP. Adenoviral CMV WT-Cre-eGFP was a gift from Dr. Kyle Schachtschneider.

All prepared constructs were validated by Sanger sequencing performed at the DNA Services (DNAS) facility within the Research Resources Center (RRC) at the University of Illinois Chicago (UIC). The relevant amino acid sequence information and plasmid resources are provided in Table 4, 5.

### Cell culture and transfection

HEK293T cells were purchased from ATCC (CRL-3216). The cells were grown in Dulbecco’s Modified Eagle medium (DMEM; Sigma-Aldrich; St. Louis, MO, USA) supplemented with 5% FBS and 1% Glutamine to 70% confluency on a plastic dish coated with 1:200 poly-L-lysine and were transfected with jetOPTIMUS® (Polyplus-transfection S.A, Illkirch, France) transfection reagent following manufacturer recommendations. The ratio of any photoactivatable Cre recombinase to the reporter plasmid in all co-transfection experiments were maintained at 1:9 and the cells were kept in the dark under aluminum foil wrap inside the incubator with 5% CO2 at 37 °C. This condition was maintained for the entire course of the experiment in the dark controls and the light activated groups except the illumination window.

### Global photostimulation with LED panel *in vitro*

24 hours post transfection, the HEK293T cells were taken out of the foil covers and stimulated with blue light, using the HQRP LED Plant Grow Panel Lamp System (3 mW/cm^2^, 470 nm wavelength). To avoid photothermal toxicity, the cell plates were placed over a perforated plexiglass rack 10 cm above the LED panel and illuminated for 4 hours with 1 second ON and 1 second OFF cycle, controlled by Arduino IDE microcontroller (Arduino Uno) and power relay (IoT Relay, Digital Data Loggers INC.) system^50^. After illumination, the cells were again foil-covered and incubated in the dark overnight (∼ 20 hours) in the incubator with 5% CO2 at 37 °C, prior to analysis (Fig. 1c, Supplementary Fig. 1a).

### Fluorescence microscopy analysis of Cre activity following global Illumination

In experiments involving LED-panel-based global illumination, 20 hours post illumination, the light and dark condition HEK293T cell samples were washed in 1X PBS and stained with Hoechst dye (Hoechst 33342; 20mM, 1 µg/mL) for 10 minutes and subsequently fixed with 2% PFA and rinsed with PBS. The procedures were performed under safe red lights until fixation. Fixed HEK293T cells in PBS were imaged using an EVOS^TM^ Auto 2 Invitrogen fluorescence microscope using 20X air objective. The miRFP670, tdTomato, GFP, and Hoechst signals were captured via epifluorescence wide-field imaging. Cre activity was measured by calculating the ratio of the number of GFP-expressing cells to the number of tdTomato-expressing cells, reflecting the conversion of the tdTomato reporter to GFP. Image processing and analysis were performed using NIH ImageJ/FiJi open-source software^85^ (Supplementary Fig. 1a).

### Flow cytometry analysis of Cre activity following global Illumination

In experiments involving LED-panel-based global illumination, 20 hours after illumination, the light and dark condition HEK293T samples were washed with 1X PBS, trypsinized with 0.25% Trypsin EDTA, resuspended in FACS buffer (1% BSA, 2 mM EDTA, 0.1% sodium azide) and filtered through 70 μm disposable cell strainers to make single cell suspensions. For Flow cytometry, the cell suspensions were run on CytoFLEX S Flow Cytometer (Beckman) and data analysis was done using Kaluza Analysis 2.1 (Beckman) software. The data analysis was performed using a gating strategy applied as the percentage of miRFP670^+^, tdTomato^+^, GFP^+^ cells. It is worth mentioning that in a 24-hour experiment, we observe no miRFP670^+^/GFP^+^ cells, that are tdTomato^−^.

### Optoelectronic device design and fabrication

Supplementary Figure 2 demonstrates the fabrication procedure of the optoelectronic device. Fabrication of the flexible printed circuit board (fPCB) began with laser ablation of a sheet of PI/Adhesive/Cu/PI/Cu/Adhesive/PI (12.5/15/12/25/12/15/12.5 μm; PCBway) to pattern both sides with circuit traces, bonding pads, and coil antennas. A separate fPCB with an identical stack-up formed the serpentine interconnect. The fabrication and assembly of the devices began with the soldering of electronic components using a soldering iron and heat air gun at approximately 350 °F. A low-temperature soldering paste (CHIPQUIK, TS391AX10) bonded the various electronic components to the fPCB. A UV-curing epoxy (UV10TKMed, MasterBond Inc.) coated the contact pads and µ-ILED (length: 270 µm, width: 220 µm, height: 50 μm, TR2227, Cree Inc.) at the tip to reinforce the solder connections. A uniform layer of PDMS (10:1 elastomer to curing agent; Sylgard 184, Dow Corning Inc.) encapsulated all circuit components (thickness: 300 µm) on the coil to form a uniform interface between the device and surrounding tissues in physiological conditions. Chemical vapor deposition formed a uniform coating of parylene-C (thickness: 10 µm; Specialty Coating Systems Inc.) on the assembled device. Polymethylmethacrylate (PMMA A5; MicroChem Inc; 2000 rpm) was spin-coated for 60 s on a silicon (Si) wafer and cured at 180 °C for 1 min on a hot plate to form a substrate. Next, a silicone layer of Ecoflex 00-30 A and B (Smooth-On Inc., 11:11 weight ratio) mixed with Sylgard 184 (4:0.4 elastomer to curing agent weight ratio) in a 4:1 weight ratio was spin-coated onto the substrate to a thickness of 300 µm. The prepared devices were immersed into the uncured silicon to completely encapsulate them, followed by curing at 100 °C for 15 minutes. Finally, a customized die was used to uniformly cut out the devices.

### Wireless operation of the optoelectronic devices

A commercial RF system (Neurolux Inc.) wirelessly delivered power to the implanted optoelectronic devices for localized optogenetic stimulation of lung tissue. The system consisted of the following components: (1) a laptop equipped with customized NeuroLux software to manage device operation, (2) a Power Distribution Control box for providing wireless power and communication to the devices, (3) an antenna tuner box designed to optimize power transfer and match the impedance between the source and the antenna, and (4) a cage box with adjustable double-loop designs to facilitate device operation in various environments (e.g., *in vitro* and *in vivo* conditions) (Supplementary Fig. 6c).

### Electrical validations in benchtop testing and modeling

A transmitting circular coil made of copper wire (diameter: 15 mm) connected to a vector network analyzer (E5063A, Keysight Technologies Inc.) measured the S11 signal of the device under bending of the receiver coil up to a radius of 2 mm. The Q-factor was calculated as the ratio between the resonant frequency (*fr*) and the full width at half-maximum (FWHM: *Δf*).

The numerical model for electromagnetic validations of the device relied on finite element analysis (Ansys HFSS, ANSYS, Inc.) to simulate the electromagnetic response as the coil is bent up to a radius of 2 mm. A matching capacitor tunes the coil to a resonant frequency; then, geometrical, resistive, and inductive changes caused by bending alter the resonance frequency, the quality factor of the coil, and the magnitude of the scattering parameter. A frequency sweep was performed between 10–18 MHz to capture the electromagnetic response and frequency shift as a function of bending radius from 10 mm to 2 mm. In the mechanics simulation, the elastomeric encapsulation was meshed with solid elements (C3D8R), and the Cu layer was meshed with shell elements (S4R). The total number of elements in the model is ∼350,000 to ensure accuracy. In the electromagnetic simulation, an adaptive meshing strategy was adopted to ensure mesh convergence over a spherical radiation boundary with a radius of 500 mm. The elastic modulus and Poisson’s ratio values used in the simulation were 119 GPa, 0.34 for Cu, and 60 kPa, 0.49 for Ecoflex. The relative permittivity, relative permeability, and conductivity of Cu used in the simulation were 1, 0.999, and 5.8 × 10^7^ S m^−1^, respectively.

### Mechanical validations in benchtop testing and modeling

The experimental analysis of the mechanical durability of the devices involved tensile and cyclic tests to determine the transition threshold from elastic to plastic deformation. A tensile tester (Tensile Tester, Mark-10 Inc.) measured the stress-strain curves of the serpentine in tension mode, with the sample clamped between the top and bottom of the cantilever and both ends of the serpentine connected to a multimeter to measure electrical continuity. The tensile modulus was calculated by fitting the tangent of the linear elastic region in the stress-strain curve. The cyclic test, using a custom-built stretching stage based on a screw-driven linear stage (ATS100, Aerotech, Inc.), applied dynamic mechanical stress to the devices, simulating a full range of natural animal movements: from −40% bending to +50% stretching for a total of 10,000 consecutive cycles at 1 Hz^40^. Sample preparation for cyclic testing involved attaching external Cu wires (diameter: 0.1 mm) to the contact pads of the devices using a soldering iron with low-temperature solder paste (Chip Quik Inc.) and a flux pen (SRA Soldering Products) to measure the resistance across the device. A block of PDMS (10:1 elastomer to curing agent; Sylgard 184, Dow Corning Inc.) was placed on the top and bottom of the device to protect it from the metal clamps during sample fixation and cyclic testing. To simulate physiological conditions in animal models, the devices were immersed in a phosphate-buffered saline solution maintained at body temperature (around 37 °C) and a neutral pH of 7.4. Measurements of the electrical resistance across the device before and after these mechanical validations identified any shift from elastic to plastic deformation. Here, no change in electrical resistance indicated elastic deformation, while a significant increase in resistance indicated plastic deformation. The numerical modeling of the mechanical durability of the devices relied on the 3D finite-element simulation (ABAQUS, Analysis User’s Manual 2023) to determine the elastic characteristics of the serpentine and coil when subjected to stretching and bending deformations. The strain level at the serpentine copper (Cu) layers was used to identify the elastic limits during deformation.

### Modeled optical properties of murine lung tissue

The absorption (*α*_0_) and scattering (*α_s_*′) coefficients of murine lung tissue at 470nm were extrapolated using an exponential fit for absorption (*k_abs_*) and a power-law fit for scattering (*a*, *b*) based on 80% inflated lung measurements at 535 nm and 730 nm^86^. The explicit equations are given as *α*_0_(*λ*) = *α*_0,*ref*_ *e^kabs^* (λref – λ) and *α_s_*′(*λ*) = *aλ^b^* , which resulted in estimates of *α*_0_ = 2.2 cm⁻¹ and *α_s_*′ = 135 cm⁻¹, respectively. Under the assumption of predominant forward scattering and a volumetrically weighted anisotropy factor for alveoli and intrinsic tissue taken as *g_a_*= 0.6 and *g_t_*=0.85^2^, the effective anisotropy factor at 80% inflation was estimated as *g* = 0.65. The Radiative Transfer Equation was used to estimate an effective attenuation coefficient of *α_eff_* = 30.3 cm⁻¹ through the equation 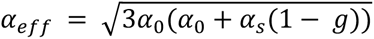^87^, where the total scattering factor is defined as 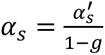^88^. The index of refraction was assumed to be 1.373 based on deflated measurements in rat lung tissue^89^. Although, it is important to note that the ∼55 µm diameter alveoli sacks in mice^90^ are modelled as air-filled microspheres and accounted for in the estimated scattering factor at 80% inflation^86,91^.

### µ-ILED emission profile modeling

Power was transferred wirelessly to the device through a near field magnetic link between a NeuroLux transmitter coil and an integrated parallel resonant receiver coil tuned to the drive frequency of 13.56 MHz^38^. The harvested voltage was rectified and then regulated to 2.8V using a low dropout regulator to drive the µ-ILED. Three potential current limiting resistors Rs = 200 Ω (580 µA), 2k Ω (135 µA), 10kΩ (37 µA) were selected to establish an appropriate illumination intensity for generating adequate physiological responses while minimizing adverse tissue effects. The near-field irradiance profiles were subsequently generated using a ray tracing approach and modelled as a Gaussian beam with a width waist of 2*W*_0*x*_ = 220 µm and 2*W*_0*y*_ = 270 µm. The model assumed transmission from a *μ*-LED encapsulated in a biocompatible epoxy (Master Bond, UV10TKMed) and coated with a thin 12 µm layer of water-proofing parylene-C into a superstrate composed of a soft 200µm interfacial layer (Ecoflex 00-30) and murine lung tissue^92^. A superstrate of 1mm^3^ within the interfacial-tissue layer was discretized into voxels with 5µm side lengths and the irradiance *I_em_*(*x*, *y*, *z*) at each voxel was calculated by scaling the normalized gaussian equation^92^

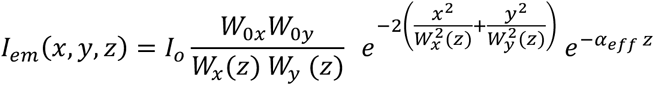

with 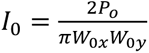, where *P* was the total measured optical power of the *μ*-LED with epoxy and parylene. The beam divergence in the x and y directions was encoded in the variables *W_x_*_,*y*_(*z*) through the equation *W_x_*_,*y*_(*z*) = *W*_0*x*,*y*_ + *ztan*(*θ_div_*_/2_), respectively^93^. Additionally, the divergence half-angle *θ_div_*_/2_ was calculated as 32.5*°* using Snell’s Law where the indices of refraction of the epoxy and Ecoflex were 1.55 and 1.405 respectively, resulting in a numerical aperture of 0.833. The effective attenuation in Ecoflex was measured^94^ to be *α_eff_* = 2.4 cm^−1^, thus this region was dominated by dispersion.

### *In vitro* photostimulation with µ-ILED and fluorescent microscopy analysis

To analyze local regulation of E-LightR-Cre by µ-ILEDs, 70% confluent HEK293T cells were co-transfected with E-LightR-Cre-miRFP670 and Floxed-STOP-mCherry reporter DNA constructs at 1:9 ratio in 35 mm tissue culture plates and incubated in the dark for 16-18 hours. Following the incubation, the 35 mm dishes were illuminated with wireless µ-ILED devices attached beneath each plate with the illuminating tip at the center (Fig. 3a). We used the Low_power µ-ILED (2 kΩ resistance, 30 µW power, 470 nm wavelength) and Ultra-low_power µ-ILED (10 kΩ resistance, 6 µW power, 470 nm wavelength) devices with 20% duty cycle, under 10 Watt (W) transmitting antenna power and 10 Hz frequency of illumination and varying illumination duration (2 hours, 6 hours or 24 hours). Illumination was carried in the RF chamber to mimic the conditions for subsequent *in vivo* experiments. A sterile wireless RF chamber (L X W X H :: 30.5 cm X 25.4 cm X 15.2 cm) was placed within the cell culture incubator to enable wireless connection. The cell culture dishes with the µ-ILED devices attached beneath, were positioned inside the chamber, to activate the µ-ILEDs. The entire setup was handled under safe red light to prevent unintended non-specific activation. Following illumination, the plates were wrapped in aluminum foil and incubated at 37 °C and 5% CO2 for additional time periods (22 hours, 18 hours or 0 hours respectively) to complete the total of 24 hours of incubation from the start of illumination (Fig. 3a-c).

At the end-point of each experiment involving µ-ILED-induced local illumination (Fig. 3a-c), the 35 mm dishes of HEK293T cells in growth media (DMEM with 5% FBS and 1% Glutamine) were transferred to the microscope imaging chamber under safe red lights and imaged using horizontal 2-dimensional area scanning using an Olympus IX83 microscope equipped with motorized Olympus SSU ultrasonic stage and the cellVivo Incubation System (Evident Scientific) that maintains the cells at 37 °C and 5% CO2 during imaging. Imaging was performed with an EMCCD camera (Andor iXon Ultra 888; Oxford Instruments) paired with a 4X air objective (UPlanXApo 4X/0.16 NA). The miRFP670 and mCherry signals were captured via confocal imaging using excitation laser lines of 647 nm and 561 nm wavelengths respectively. Image acquisition was performed using cellSens software (version 4.1), enabling automated alignment of multiple images with a 10% overlap. Each cell plate imaging took approximately 10 minutes. The extent of localized Cre activity was determined by measuring the radius of mCherry reporter expression on each plate. For image processing, the stitching artifact was removed using Fast Fourier Transformation (FFT) in ImageJ ^85^. After that, the diameter of the activated zone was determined using the image processing functionalities in MATLAB (version R2022b). Each image was thresholded for removal of background signal and the approximate center of the illuminated region was marked. Subsequently, two rectangular regions (400×60 pixels and 60×400 pixels, respectively) were cropped from the binarized mCherry signal distributions. The corresponding pixel values, represented as 2D matrices, were averaged along their shorter dimension to produce two 1D vectors, each 400 elements long, representing the average intensity distributions along the horizontal (x) and vertical (y) directions in the image. The average intensity distributions I(x) and I(y) were then fitted to Gaussian curves as:

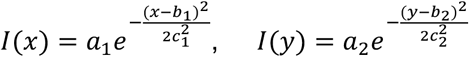

where, *a* is the height of the curve’s peak, *b* the position of the center of the peak, *c* the standard deviation, and subscripts 1 and 2 represent fit parameters corresponding to x and y directions, respectively. After that, the average radius of the activated zone was calculated as *r* = (2*c*_1_ + 2*c*_2_)/2, where the half-width (radius) of the activated zone in each direction was assumed to be twice the respective standard deviation, which corresponds to approximately 95.5 % of the total area under the Gaussian curve^95^ .

### Local expression of oncogenic KRas and analysis by fluorescence-activated cell sorting and bulk RNA sequencing

To stimulate localized expression of GFP-KRas(WT/G12D) constructs, 70% confluent HEK293T cells were co-transfected with E-LightR-Cre-miRFP670 and one of the three constructs: pCA-mTmG(GFP-only), pCA-mTmG(GFP-KRas-WT), or pCA-mTmG(GFP-KRas-G12D) in 1:9 ratio and incubated in dark for 16-18 hours. Following incubation, local illumination was performed using Low_power µ-ILED devices operating at 10 W power, 10 Hz frequency, and 20% duty cycle for 6 hours. Following illumination, the plates were wrapped in aluminum foil and incubated at 37°C and 5% CO2 for 18 hours in the dark. At the end points, the cells were harvested under safe red light for fluorescence-activated cell sorting (FACS) of GFP positive and GFP negative population (Fig. 3d).

The cells were harvested as single cell suspensions and the GFP-positive and GFP-negative cells from these locally illuminated plates were sorted using the MoFlo Astrios (Beckman Coulter). The sorted cells were used for total RNA extraction by TRIzol Reagent (Invitrogen, 15596026), the RNA was treated with DNase, and its quality was evaluated with gel QC. All handling of the live cells was performed under safe red light, to avoid unwanted non-specific activity. Bulk RNA sequencing was performed with oligo-dT mRNA directional using Illumina Novaseq X plus for coding genes at the University of Illinois Urbana-Champaign Genomic Facility.

### Bulk RNA sequencing analysis

A reference transcriptome index was created using Gencode hg38 annotations v45 to map against. To distinguish transgenic KRasG12D transcripts from endogenous KRas if any, the KRasG12D sequence was added to the reference, as even in the absence of DNA recombination the reporter construct design still leads to transcription of a single mRNA encoding tdTomato, GFP, and KRas-G12D genes. However, KRas-G12D protein is not produced due to the stop codon at the end of tdTomato gene. Sequenced reads were mapped to the reference genome (GRCh38) and their abundances were quantified using Salmon v1.3.0 in quasi-mapping mode^96^. Salmon produced transcript-level quantification files, which were subsequently aggregated into gene-level counts for downstream analysis. Differential expression analysis was performed using DESeq2 v1.40.2, which models count data based on the negative binomial distribution, suitable for identifying differentially expressed genes across conditions^97^. Normalization and dispersion estimation were automatically performed by DESeq2 to account for library size differences and biological variability. Genes with an adjusted p-value < 0.05 were considered statistically significant. Differentially expressed genes were further analyzed for enrichment in Gene Ontology (GO) Biological Processes using the clusterProfiler package v4.8.2^98^ .

### Animal model

All animal experiments were performed under the protocol approved by the office of Animal Care and Institutional Biosafety (OACIB) at the UIC. Mice were maintained in a pathogen-free environment at the University of Illinois Animal Care Facility with 12 hours dark and 12 hours light cycle, temperature at approximately 23 °C, and humidity at 40-60%. Mice had free access to water and food. For local optogenetics experiments (Fig. 5b, Supplementary Fig 7a); 8-10-week-old male or female Rosa-LSL-tdTomato mice (B6.Cg-*Gt(ROSA)26Sor^tm^*^9^(CAG–TdTomato)*^Hze^*/J; The Jackson Laboratory strain #:007909)^65^ or Rosa-LSL-tdTomato/ SPC-GFP (obtained from Dr. Yuru Liu; UIC) mice were used ^99^. For tumor initiation studies (Fig. 5c-f; Supplementary Fig. 8a-c), 8-12-week-old male and female LSL-Kras-G12D strain mice (LSL-K-ras G12D/B6.129S4-Kras^tm4Tyj^/J strain; The Jackson Laboratory Strain #:008179) were used^19^.

### Wireless µ-ILED surgical implantation

The animals were anesthetized with Ketamine/Xylazine (100/5 mg/kg body weight) administered intraperitoneally. Once anesthesia was confirmed, Buprenorphine ER (0.3 mg/kg) was delivered subcutaneously, and the surgical procedure was followed. (1) The trachea was visualized using a surgical microscope (Amscope MU500-HS) to ensure proper placement of the catheter for ventilation. (2) The animal was intubated with a 20G catheter and placed under isoflurane anesthesia on an isothermally heated surgical stage set at 37 °C and ventilated with a VentElite Small Animal Ventilator (Harvard Apparatus, Holliston, MA) (120 bpm, 14 PIP, and a flow volume of 500-700 µL). (3) The posterolateral skin area over the ribs was shaved and cleaned with an alcohol swab, and a vertical 1 cm incision was made on the skin over the 5^th^ rib. (4) The incision was extended layer by layer through the muscles to expose the ribs and the ventilated lung beneath. (5) A small vertical incision was made in the intercostal muscle between the 6^th^ and the 7^th^ ribs to insert the device tip with the µ -ILED (∼3 mm wide) facing the lung. (6) The entire surgical stage was rotated 90 ° for superior surgical access. A reverse cutting P-3 needle (13 mm, 3/8) with a 6-0 silk suture (#S-N610R13-B; AD Surgical) was passed through the 5^th^ and 6^th^ rib, staying closer to the 5^th^ rib, while avoiding the lung underneath and exiting the thoracic cavity through the incision space between the 6^th^ and 7^th^ ribs. (7) The needle was then passed through the tip of the optogenetic device, through the Ecoflex-parylene coating, next to the μ-ILED tip, to create a loop, ensuring the device is oriented with the receiver coil side facing upward, μ-ILED side facing downward and entering from the top and exiting at the bottom. (8) The needle was reintroduced into the chest cavity, through the incision space between the 6^th^ and 7^th^ ribs and made to exit through the intercostal muscle between the 4^th^ and the 5^th^ ribs, staying close to the 5^th^ rib. (9) While holding both ends of the suture, the device was slowly inserted into the chest cavity. (10) Both ends of the suture were carefully pulled further so that the device was positioned underneath the ribs. (11) The two ends of the suture were tied around the 5^th^ rib, securely, but not too firmly, to avoid any breakage of the ribs, ensuring to secure the device in position. (12) A second needle with 6-0 silk suture was passed inside the thoracic cavity between the 7^th^ and 8^th^ rib and then exited between the 5^th^ and the 6^th^ rib to suture ribs 6 and 7 together and seal the incision. (13) Before tying off the suture, the initial knot was made without pulling it tight, to prepare for sealing the chest cavity. (14) The suture was then tied to completely seal the chest cavity, following an inspiration hold and a gentle squeezing of the chest to make sure to remove the excess air out of the chest cavity. (15) After resealing the thoracic cavity, the serpentine interconnect was extended posterolaterally through the muscle layers underneath the skin with the help of blunt end forceps creating a cavity right under the skin layer for the device to slide into. (16) This ensured subcutaneous positioning of the NFC enabled receiver coil and the attached red indicator LED on the back of the mouse for optimal power harvesting and operational monitoring. (17) The surgical incision in the muscle and skin layers were closed with 5-6 interrupted sutures (6-0 nylon sutures with reverse cutting P-3 needle: 13 mm, 3/8). Halfway through the skin sutures, the isoflurane flow was shut off, to gradually allow the animal to come back to consciousness. (18) It was ensured that the incision is securely sealed by all sutures. The animals were then disconnected from the ventilator, and the intubation tube was removed. Once normal spontaneous breathing resumed, the mice were allowed to recover on an isothermal pad with access to food and water. Appropriate postoperative care was provided for a minimum of three days following surgery. The animals were observed every week following implantation, until the end of the experiment. The numbers in the text represent the steps shown in the Supplementary Fig. 6a. Extended surgical procedure will be found in Supplementary File S1. All procedures were conducted according to the protocols approved by the Institutional Animal Care and Use Committee at UIC.

### Micro-CT

Mice with implanted µ-ILED devices were anesthetized with 1-2% isoflurane in oxygen (O2) during imaging. A Bruker preclinical x-ray micro-CT imaging system (Skyscan 1276, Bruker Belgium, Kontich, Belgium) was used to acquire the CT images. First, respiratory gated scans were acquired of the lungs with the following parameters: 4 × 4 binning, 65 kV/200 µA, 0.25mm Al filter, 160 ms exposure, 37 µm voxel size, 3600 projections over 360 °, and two frame averages. Images were retrospectively sorted into 4 bins for respiratory gating and reconstructed with filtered backprojection using NRecon v2.2.0.6 (Bruker Belgium, Kontich, Belgium). Then, the mice were overdosed with isoflurane until the respiratory monitor no longer registered any breathing. The whole-body, postmortem scans were acquired with the following parameters: 2 × 2 binning, 85 kV/200 µA, 0.5 mm Al filter, 150 ms exposure, 18.5 µm voxel size, 3600 projections over 360 °, and two frame averages. Filtered backprojection reconstruction was used without gating. Amira 2024.2 (FEI Co, Hilsboro, OR) was used to merge the gated lung CT images with whole-body postmortem scans. Then the device, lungs, and skeleton were segmented, followed by 3D surface rendering. For these procedures, the device-implanted mice were transferred from the UIC to the Northwestern University (NU), by an approved, specialized courier. All procedures were approved by the animal institutional care and use committees at UIC and NU.

### Behavioral experiments

The animals were placed in a 30 × 18 -cm arena. Each mouse was recorded individually for approximately 25 minutes using a smartphone camera (iPhone 13) mounted on a tripod, capturing video at 1080p resolution (1080 × 1920) and 25 frames per second (FPS). Mouse movement was tracked using the Track-Analyzer module in the Animal Tracker API plugin in ImageJ^100^. Briefly, the image sequence was obtained from an open-source MP4-to-JPEG converter and imported into ImageJ. A polygonal area was selected on the image to define the analysis boundary, detected by the Radial Maze selection under the Animal Tracker plugin, generating a tracking window. Background subtraction was performed using Gaussian Blur, and the mouse was thresholded in the processing window. Analysis started at frame 7501 (5 minutes) and ended at frame 30000 (20 minutes), including a 5-minute acclimatization period at the beginning. The thresholded mouse contour was selected to initiate tracking from the Tracker menu. After tracking, analysis zones were selected from the Show Analyzer menu, and the correct frame interval and pixel/unit values were entered to generate raw data for distance and velocity vectors. The data were entered into Microsoft Excel and analyzed for consecutive 5 minutes time points to measure the average distance traveled and average velocity from three 5-minute intervals per video. The detailed protocol is available in Supplementary File S2.

### Adenoviral Cre delivery

The mice received a global lung targeted endotracheal delivery of 5X10^8^ PFU (in ∼30 µl) of Adenoviral CMV-E-LightR-Cre-miRFP670 or unmodified Adenoviral CMV-WT-Cre-GFP. For oropharyngeal endotracheal intubation and Adenoviral Cre delivery to the lung, the mice were anesthetized with Ketamine/Xylazine (100/5 mg/kg body weight) administered intraperitoneally. Then they were laid on an intubation stand (ETI-MSE-01; Kent Scientific) suspended at a 45° angle and held in position with surgical thread secured around their incisors, and the tail immobilized with adhesive tape. A Stereo microscope (SM-3TZ-54S-5M; Amscope) with LED-intensity adjustable ring light (LED-144W-ZK; Amscope) was used to allow illumination and magnification of the larynx inside oral cavity. In the next step, the tongue was gently pulled out and held with the recessive hand to visualize the larynx under the microscope. Once the epiglottis and the arytenoid cartilages were visualized, the dominant hand was used to carefully insert a 20 G, 1.25 inch catheter over the epiglottis and between the arytenoid cartilages and gently pushed ∼5 mm deeper through the trachea. Once the catheter was placed, the intubated mice with the surgical stand were transferred into the biological safety cabinet for endotracheal delivery of the adenoviral load. Administration of the payload inside the catheter automatically delivers it through the catheter to the lungs. The catheter was removed immediately after delivery and the mice were restored from the intubation rack to be put back in their cages with access to food and water. After adenoviral delivery, the mice were maintained following the Institutional Biosafety Committee’s guidelines at UIC.

### *In vivo* illumination using implanted µ-ILED devices

To induce local optogenetic E-LightR-Cre activity in mouse lungs *in vivo*, implanted Low_power µ-ILED (2 kΩ resistance, 30 µW power, 470 nm wavelength) and Ultra-low_power µ-ILED (10 kΩ resistance, 6 µW power, 470 nm wavelength) devices were used, to deliver illumination under 10 W transmitting antenna power and 10 Hz frequency with duty cycles ranging from 20% to 50% in the RF cage. The mice were housed in a standard animal cage that was positioned within a wireless RF chamber (L X W X H :: 30.5 cm X 25.4 cm X 15.2 cm) to enable successful wireless operation and illumination on the left lung with the implanted µ-ILED. The indicator red LED on the back ensured monitoring of successful wireless connection and desired pulsing (Supplementary Fig. 6c).

### Lung tissue preparation

Mice were sacrificed at the designated time points for each experiment following approved Animal Care Committee (ACC) guidelines at UIC. The chest cavity was opened using scissors to expose the heart. The right atrium was then cut, and the heart was slowly perfused through the left ventricle, with 10 mL of PBS through a 21G needle. Thereafter, the animals were intratracheally perfused with 10 mL of PBS through a 21G needle. The perfusion was repeated with 5 ml each of 4% PFA in PBS, pH 7.4; transcardially and intratracheally. While the lungs were still being perfused with 4% PFA intratracheally, the trachea was tied with a suture to maintain the lungs in an inflated condition. The lungs were carefully removed from the thoracic cavity, with particular care taken to avoid damaging the lung, which had an implanted µ-ILED device positioned adjacent to the organ and still attached within the thoracic cavity. The extracted lungs were fixed overnight in 4% PFA at 4 °C, and each lung lobe was separated from each other the next day, prior to the subsequent analysis.

### Lung tissue clearing

To evaluate local gene recombination following local activation of E-LightR-Cre, mouse lungs were cleared for subsequent Light-sheet microscopy analysis. Lung samples from Rosa-LSL-tdTomato and Rosa-LSL-tdTomato/SPC-GFP mice were collected 10 days post illumination. After 24 hours of fixation in 4% PFA at 4 °C, the lung lobes were washed 4-5 times for 10 minutes each in PBS, then incubated in RBC lysis buffer (Invitrogen, Catalogue: 50-112-9751) overnight at 4 °C and washed again in ultrapure water (Invitrogen, Catalogue: 10977015) 4-5 times for 10 minutes each. To retain the endogenous fluorescent signals, an aqueous based tissue clearing was performed following the “Fast 3D clear” protocol with some alterations in incubation windows adapted for the mouse lungs^66^. For dehydration and delipidation, the tissue was transferred to glass vials, and incubated in a tetrahydrofuran (THF) gradient: 50% THF, pH 9.0 for 1.5 hrs, 70% THF, pH 9.0 for 1.5 hrs, and 90% THF, pH 9.0 for 18 hrs. The samples were then rehydrated in reverse order followed by washing with ultrapure water 4-5 times for 10 minutes each. Finally, the samples were incubated in Fast 3D aqueous clearing solution with refractive index (RI) 1.512-1.515 for 24-48 hours at 37 °C, then stored in the clearing solution at 4°C prior to imaging. During all these steps, the incubation chambers were aluminum foil covered to protect from any light-induced signal damage.

### Light sheet fluorescence microscopy (LSFM) of cleared lungs

The cleared lung samples were imaged using Zeiss Z7 light sheet fluorescence microscope equipped with dual sCMOS cameras (PCO.edge 4.2) following detailed imaging protocol by Yu et al^67^ . ZEN 3.1 LS (black edition) imaging software was used. The imaging system was equipped with a 5X, 0.16 NA, variable refractive index (RI) detection objective and two 5X, 0.1 NA, variable RI illumination objectives, all set to RI of 1.51. The cleared lungs were mounted on the sample holder vertically with superglue (Loctite Ultragel control) and imaged in Cargille immersion oil type A with RI = 1.515 (Cargille 16842) in the sample reservoir. The specimen navigator was used to adjust the position of the camera relative to the sample. The light sheet illumination was configured for aligned dual illumination in the desired laser channel. The software suggested value for Z-plane distances, and 10% for the tile overlap were applied throughout the experiments. Reporter tdTomato was imaged using 561 nm excitation laser. SPC-GFP was imaged using 488 nm excitation laser. Autofluorescence was imaged using 638 nm excitation laser to define the lung edges and airways. For samples containing no SPC-GFP, 488 nm laser was used for background autofluorescence imaging.

### Light sheet image processing and image analysis

The original Zeiss imaging file (.czi) was converted using Imaris File Converter (Oxford Instrument, Version:10.2.1). Views were then stitched together using Imaris Stitcher (Oxford Instrument, Version: 10.2.1) with the “Automatic Align” function. Images and videos were captured and analyzed by Imaris (Oxford Instrument, Version 10.2.1). Representative 3D micrographs were generated based on thresholded fluorescence intensities, using the Volume and Surface rendering tools of Imaris software (Oxford Instruments, Version 10.2.1). Local activation was measured as the radii encompassing 95% of the total signal area.

### Histological analysis

For histological analysis of mouse lung samples, the left and right lung lobes from different experimental groups were transferred to 70% ethanol after 24 hours of fixation in 4% PFA. The samples were then processed and embedded in paraffin at the Research Histology Core, University of Illinois Chicago. 5 µm thick consecutive sections were collected from the paraffin blocks of lung samples, using a microtome (Reichert-Jung 2030 Biocut Manual Microtome). Special attention was paid during embedding and sectioning, to obtain sections close to the µ-ILED device attachment surface on the left lung. Selected sections were stained with Hematoxylin and Eosin following previously reported protocol^101^. The sections were examined blind, for histopathological annotation. For light microscopy analysis, histological images were obtained using Aperio brightfield whole slide scanner and imager using 40X (0.25 μm/pixel) objectives at Research Histology and Tissue Imaging Core at University of Illinois Chicago. The images were visualized and extracted using QuPath (Edinburgh, UK), an open-source quantitative Pathology & Bioimage Analysis software^102^ .

### Immunohistochemistry

Immunohistochemical staining for Ki-67 (Rabbit polyclonal; Abcam ab15580) was performed on selected sections following standard protocols at Histowiz, Inc. (Brooklyn, NY, USA). Digitized images obtained from Histowiz were analyzed using NIH ImageJ/FiJi open-source software to assess number of Ki67 positive nuclei in selected regions of interest (ROIs).

### Immunofluorescence

For immunofluorescence, the sections were deparaffinized in xylene and re-hydrated using graded ethanol series and distilled water. The antigen retrieval was performed by boiling the slides in 10 nM sodium citrate buffer (pH 6.0) in a 2100 Antigen Retriever instrument (Electron Microscopy Sciences, Hatfield, PA). After Antigen retrieval, the sections were washed in deionized water and PBS for 10 minutes each and blocked at room temperature (RT) for 2 hours in PBS containing 0.3% Triton X-100 and 1% bovine serum albumin (BSA). The slides were then incubated overnight at 4°C with rabbit monoclonal anti-EpCAM antibody (cell signaling technology, catalogue number: 42515S; dilution: 1:500) and rat monoclonal anti-Ki67 antibody (BioLegend; catalogue number: 151202; dilution: 1:100) diluted in blocking buffer. Following the primary antibody, the sections were washed 3 times for 5 minutes with 0.3% Triton X-100 in PBS at RT. The sections were then incubated at room temperature (RT) for 1 hour with Goat anti-Rabbit IgG (H+L) Highly Cross-Adsorbed Secondary Antibody, Alexa Fluor™ 488 (Thermo Fisher Scientific, catalog no. A-11034; dilution 1:500) and Donkey anti-Rat IgG (H+L) Cross-Adsorbed Secondary Antibody, DyLight™ 550 (Thermo Fisher Scientific, catalog no. SA5-10027; dilution 1:500), dissolved together in blocking buffer. After secondary antibody incubation, the sections were stained with Hoechst dye (Hoechst 33342; 20 mM, 1 µg/mL) in PBS for 5 minutes followed by a 5-minute PBS wash. The slides were then mounted in Fluoromount-G (SouthernBiotech; catalogue number: 0100-01). Images were captured using Olympus IX83 microscope using EMCCD camera (Andor iXon Ultra 888, Oxford Instruments) and a 40X silicone immersion oil (UPlanSApo 40X Silicone/1.25 NA) objective. Extended Focal Imaging was performed using 488 nm laser, 561 nm laser and wide-field DAPI filter set. The proportion of proliferating epithelial cells was determined by manually counting the dual-positive EpCAM and Ki67 cells in each field of view and calculating the percentage of these cells relative to the total number of Ki67-positive cells within the same field. Image analysis was performed using QuPath: open-source quantitative Pathology & Bioimage Analysis software^102^.

### Statistics

Data analyses were performed using GraphPad Prism (Version 10.2.2. for windows, GraphPad Software, Boston, Massachusetts USA). Data are presented as mean ± 95 % confidence interval (CI). Statistical details for specific experiments—including numbers of repeats (n), P values, and statistical tests are specified in the figures and the figure legends. Data were analyzed by Brown-Forsythe and Welch’s one-way ANOVA followed by post hoc Dunnett’s T3 multiple comparisons test for unpaired data from a Gaussian distribution but unequal variance. Repeated-measures one-way ANOVA with post hoc Tukey’s multiple comparisons test was used to analyze three or more paired/matched groups. When comparing between two conditions, an unpaired two-tailed t-test (data normal, equal variance) with Welch’s correction was used for statistical significance testing. Significance levels are denoted as: ns, not significant, * : p < 0.05, ** : p < 0.01, *** : p < 0.001, **** : p < 0.0001.

## Acknowledgements

This work was supported by NIH grants R33CA258012, P01HL151327 and UIC Cancer Center Team Science grant to A.V.K. and J.R., NIH grant R35GM145318 to A.V.K. We thank Dr. Preetish Kadur Murthy for valuable discussions related to the work and Dr Asrar B. Malik for his continued support. We are grateful to Dr. Tamal Roy, for discussions and technical support with MATLAB-based data processing. We thank the Research Resources Centers at University of Illinois Chicago: RRC Histology and Tissue Imaging Core, CCRV-RRC Viral Vector Core, RRC Fluorescence Imaging Core, RRC Flow Cytometry Core, UIC for the Facility resources. CT Imaging work was performed at the Northwestern University Center for Advanced Molecular Imaging (RRID:SCR_021192) generously supported by NCI CCSG P30 CA060553 awarded to the Robert H Lurie Comprehensive Cancer Center. This work used the NUFAB facility of Northwestern University’s NUANCE Center for the development of devices, which has received support from the SHyNE Resource (NSF ECCS-2025633), the IIN, and Northwestern’s MRSEC program (NSF DMR-2308691). The Querrey-Simpson Institute for Bioelectronics at Northwestern University supported this work.

## Author Contributions

A.V.K. conceived the overall idea and supervised the project; contributed to the experiment design and planning; oversaw data interpretation. T.B. designed, planned and executed most of the cell-based and animal studies unless otherwise stated; engineered E-LightR Cre; conducted molecular cloning, cell culture and photo-stimulation experiments; performed fluorescence microscopy, flow cytometry; optimized and performed *in vivo* illumination; optimized and performed lung tissue clearing, light-sheet microscopy, image processing; performed and analyzed post-operative device biocompatibility and behavioral tests in mice, conducted H&E and IF staining and histopathological evaluation; conducted overall data interpretation and all statistical analyses; guided C.C. in optimization of surgical implantation, animal care and animal line maintenance. C.C. performed all optoelectronic device implantation surgeries; was primarily responsible for animal care, animal line maintenance, genotyping, post-operative animal care; performed paraffin sectioning. T.B. and C.C. performed endotracheal adenovirus delivery and ex-vivo lung harvest. Y-M.K. performed FACS sorting, total RNA extraction and sample preparation for bulk RNA sequencing. C.R.H. performed and analyzed the micro-CT. O.D. provided blinded identification of lung histopathological status. V.S. performed analysis of Bulk RNA sequencing data, generated the data visualization plots. J.M. performed molecular cloning. S.V. assisted Y-M.K. in FACS sorting, total RNA extraction and sample preparation for bulk RNA sequencing. Y.L. generated and provided the Rosa-LSL-tdTomato/SPC-GFP mouse line and contributed to discussions on its application. P.T.T. provided guidance and technical assistance with light sheet microscopy image acquisition and image processing. S.S-Y.L. provided guidance and software resources for light sheet image processing. J.R. co-conceived the concept with A.V.K., oversaw data interpretation. T.B., and A.V.K. wrote the manuscript with inputs from J.R. C.C., Y-M.K., V.S., O.D., C.R.H., P.T.T. contributed to manuscript preparation. T.B. prepared the figures.

M-K.L., C.H.G., and J.A.R. conceptualized the design of the optoelectronic device. M-K.L. and Y.C. designed and optimized the development of devices. M-K.L., Y.C., J.K., and K.M. performed fabrication of devices. R.A., D.H. and Y.H. performed theoretical modeling of the device performance. M-K.L., K.Y., J.K., K.M., D.H., M.K., S.L., S.C., H.H., and Y.K. performed experimental validation of the device performance. M-K.L., C.H.G., and J.A.R. prepared the manuscript and figures related to device design, including theoretical simulations, experimental performance analysis and overall impact.

---

**Supplementary Fig. 1:**
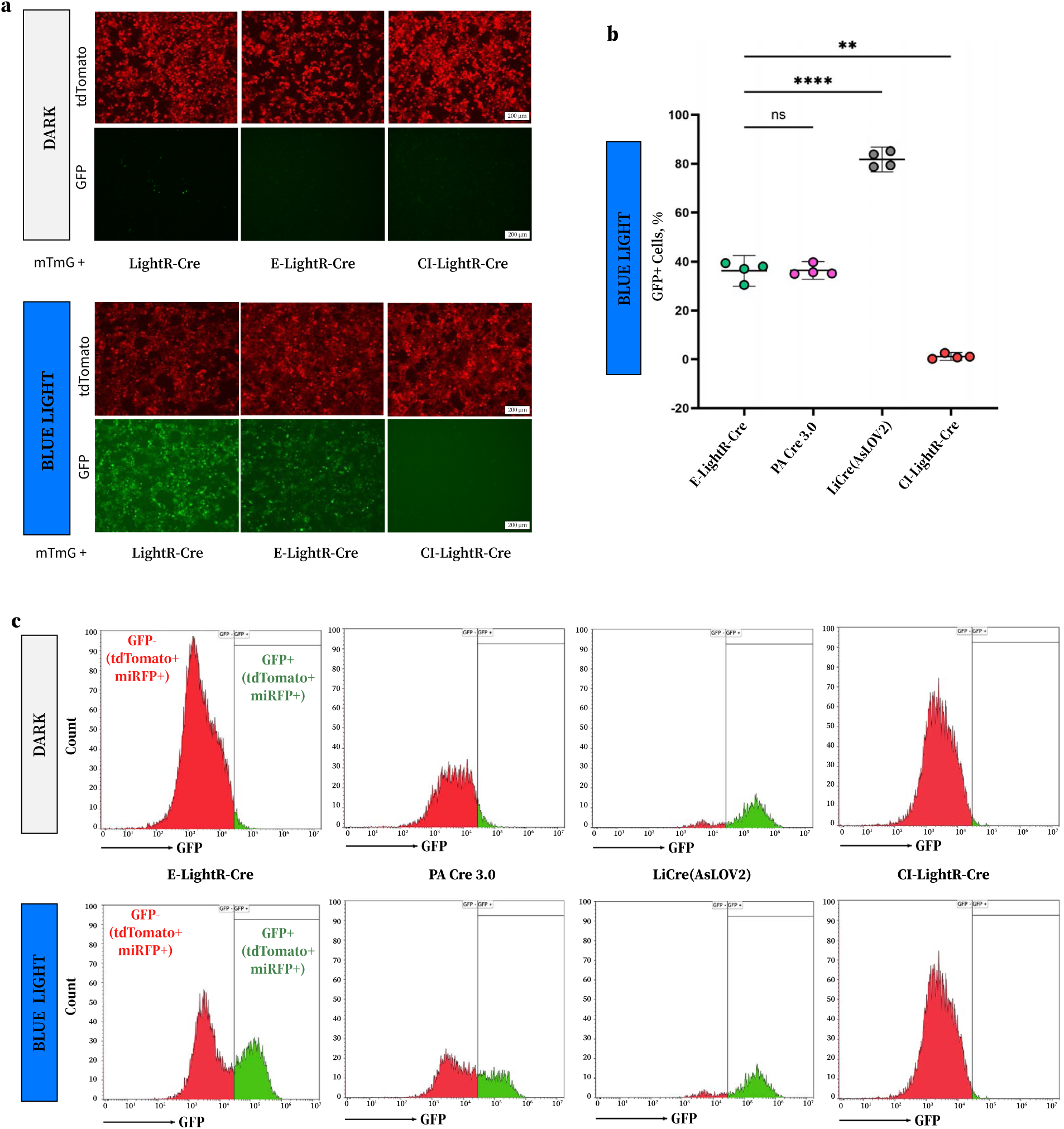
Development of light-regulated Cre recombinase demonstrating no leaky activity in the dark. **a**, Representative epifluorescence images of HEK293T cells transiently co-transfected with the indicated Cre construct and a ‘loxP-tdTomato-STOP-loxP-GFP’ reporter plasmid (pCA-mTmG) either kept in the dark or illuminated with blue light. Images were taken 20 hours following illumination. Scale bars 200 µm. **b**, Flow cytometry analysis of indicated Cre construct activity following blue light illumination. Fraction of GFP expressing cells was determined using gating strategy in **c**. Mean ± 95% confidence interval is shown for n=4 experimental replicates with statistical significance assessed by Brown-Forsythe and Welch one-way ANOVA with Dunnett’s multiple comparisons test. ns: not significant, p > 0.05; **: p < 0.01; **** : p < 0.0001. **c**, Representative histograms from flow cytometry analysis illustrating the gating criteria for selection of GFP-expressing population of cells.

**Supplementary Fig. 2:**
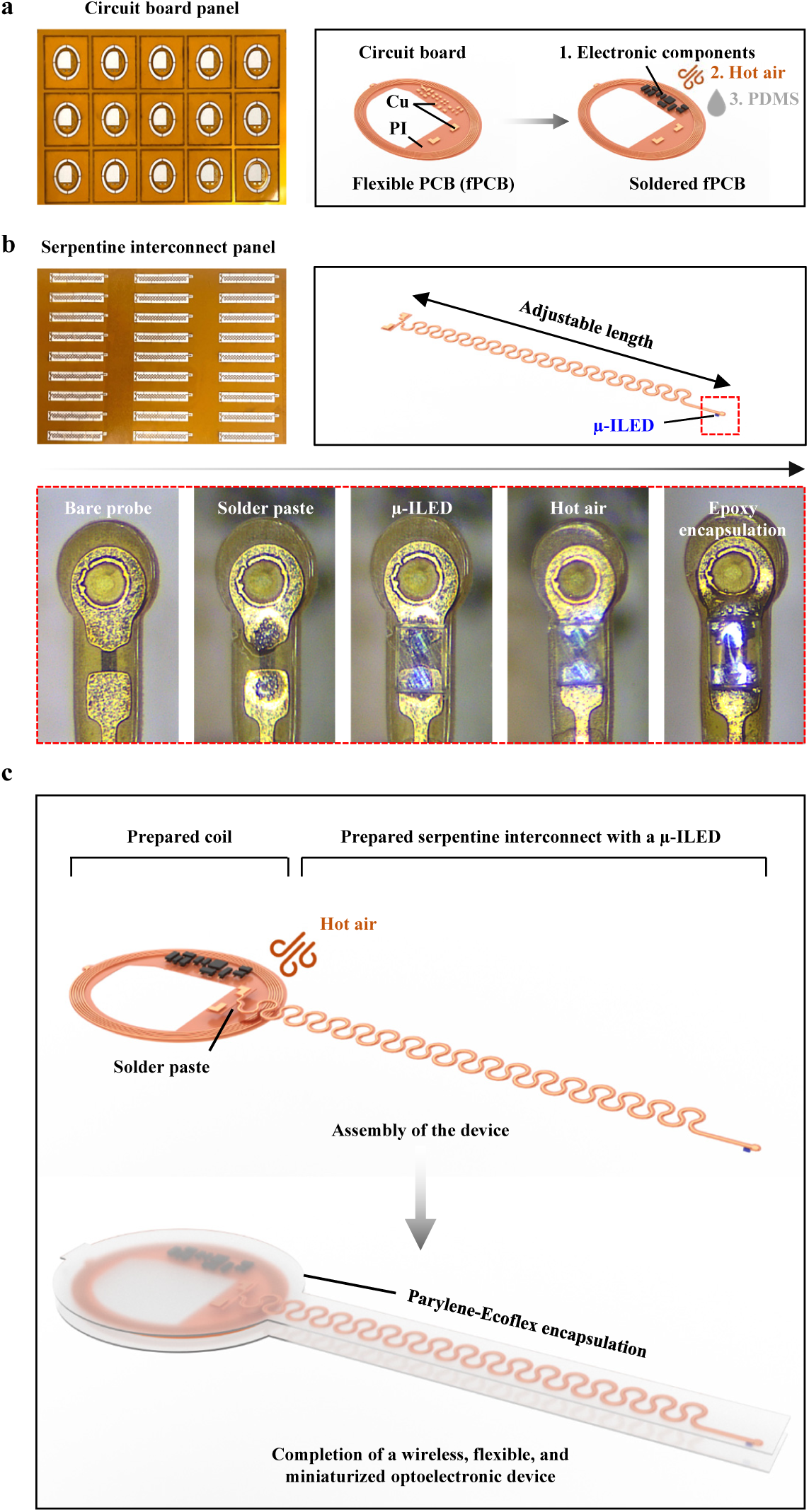
Fabrication of the optoelectronic device. **a**-**c**, Schematic illustrations of optoelectronic device fabrication; **a**, Image of the circuit board panel (Left) and soldering of electronic components onto the circuit board (Right). **b**, Image of the serpentine interconnect panel (Left), soldering of a µ-ILED at the tip (Right), and a corresponding step-by-step illustration of the soldering of µ-ILED (Bottom). **c**, Final assembly and encapsulation of the device, highlighting the completion of a wireless, flexible, and miniaturized optoelectronic device.

**Supplementary Fig. 3:**
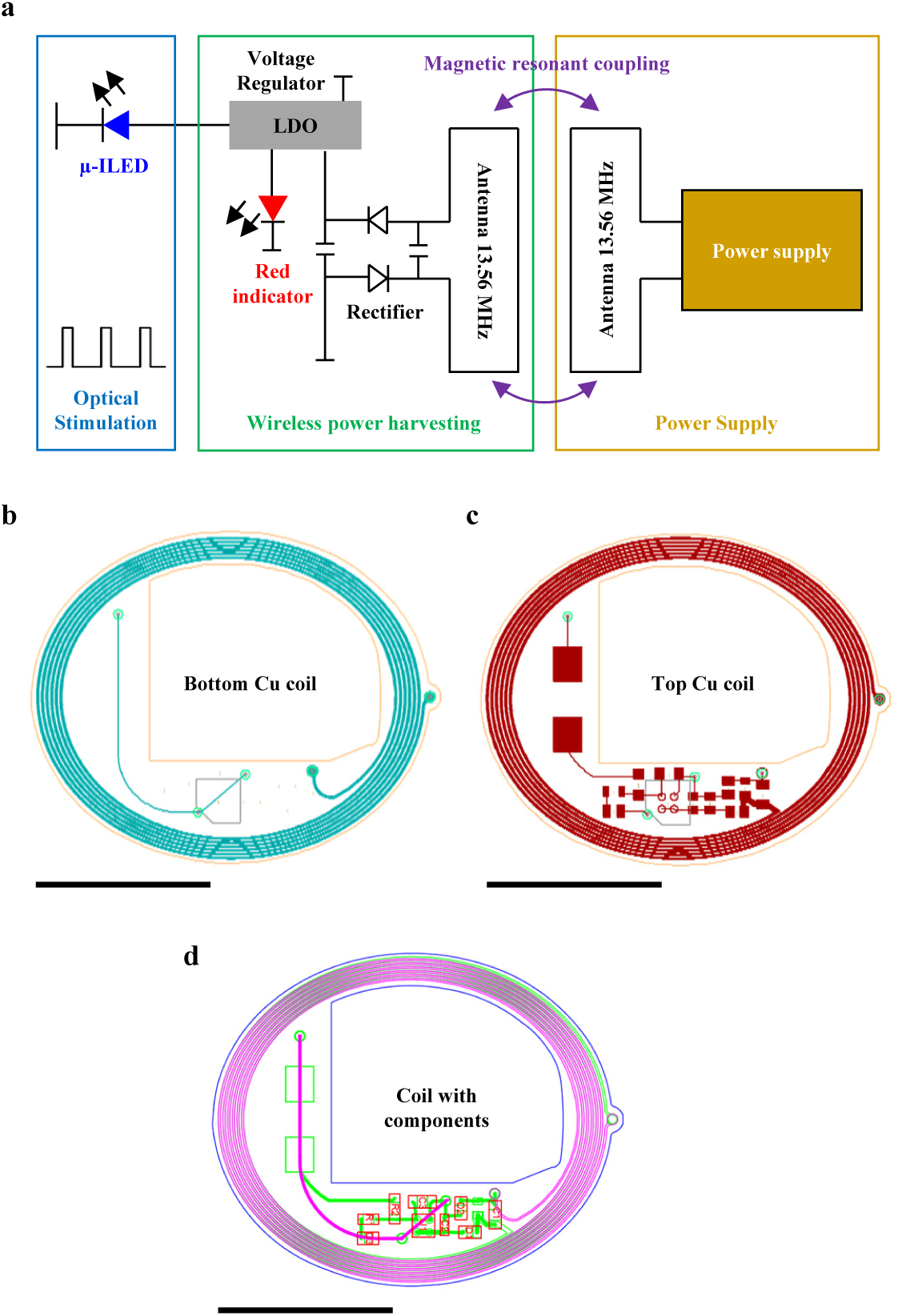
Wireless optoelectronic circuitry design. **a**-**d**, Schematic illustrations of optoelectronic device circuitry design; **a**, Circuit diagram. **b**-**d**, CAD designs of receiver coil with electrical circuitry. Scale bar 5 mm.

**Supplementary Fig. 4:**
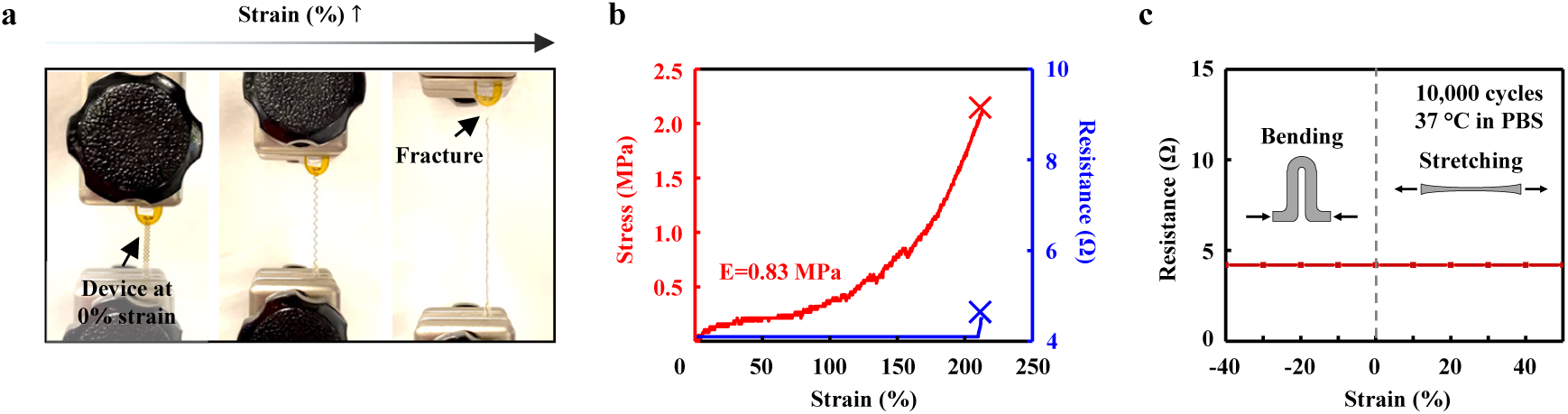
Mechanical properties and durability of the serpentine interconnect. **a**, Image of tensile test. **b**, Stress-strain curve of the serpentine traces and corresponding electrical resistance of the device measured during the tensile test. The onsets marked “X” indicates the moment of interconnect fracture. **c**, Repetitive cyclic testing of the flexible and stretchable interconnects used in the optoelectronic device, showing changes in electrical resistance over 10,000 cycles of stretching and compression at 1 Hz under physiological conditions. The stable resistance (red line) indicates the mechanical durability of the device under mechanical deformation without mechanical fracture.

**Supplementary Fig. 5:**
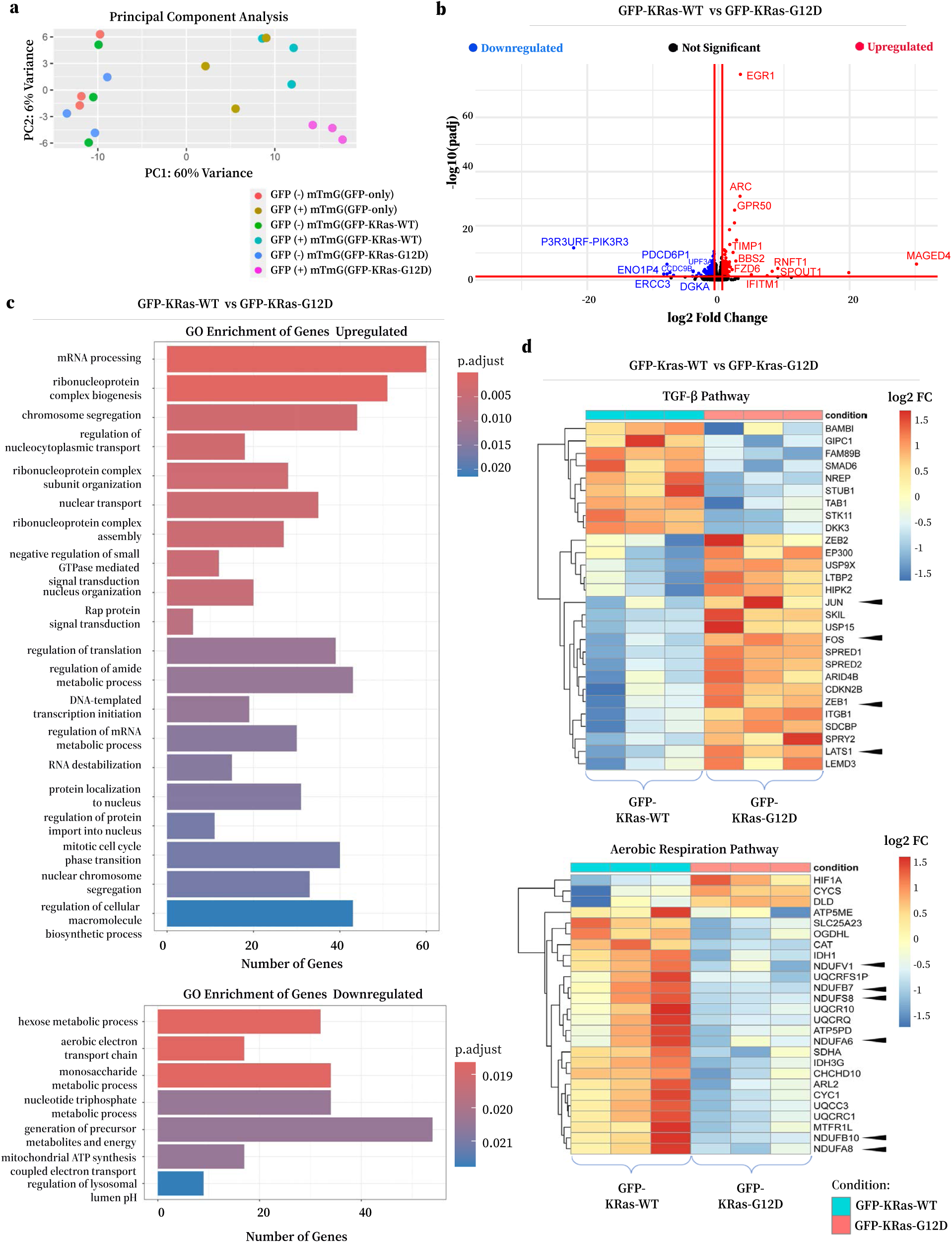
Characterization of E-LightR-Cre mediated spatiotemporal control of KRas-G12D expression *in vitro*. **a**, Principal component analyses (PCA) plots of RNA-seq data showing indicated groups of cells. **b**, Volcano plot of the top differentially expressed genes comparing GFP-KRas-WT to GFP-KRas-G12D expressing cells. Top 17 differentially expressed genes are annotated. **c**, Gene Ontology (GO) enrichment analysis of differentially upregulated (Top) and downregulated (Bottom) pathways comparing GFP-KRas-WT to GFP-KRas-G12D expressing cells. Significance criteria used: adj. p-value < 0.05. **d**, Heatmaps of differentially expressed genes in TGF-β Pathway (Top) and Aerobic Respiration Pathway (Bottom). Significance criteria: adj. p-value < 0.05. Color-coded scale corresponds to indicated fold change in gene expression (log2 FC).

**Supplementary Fig. 6:**
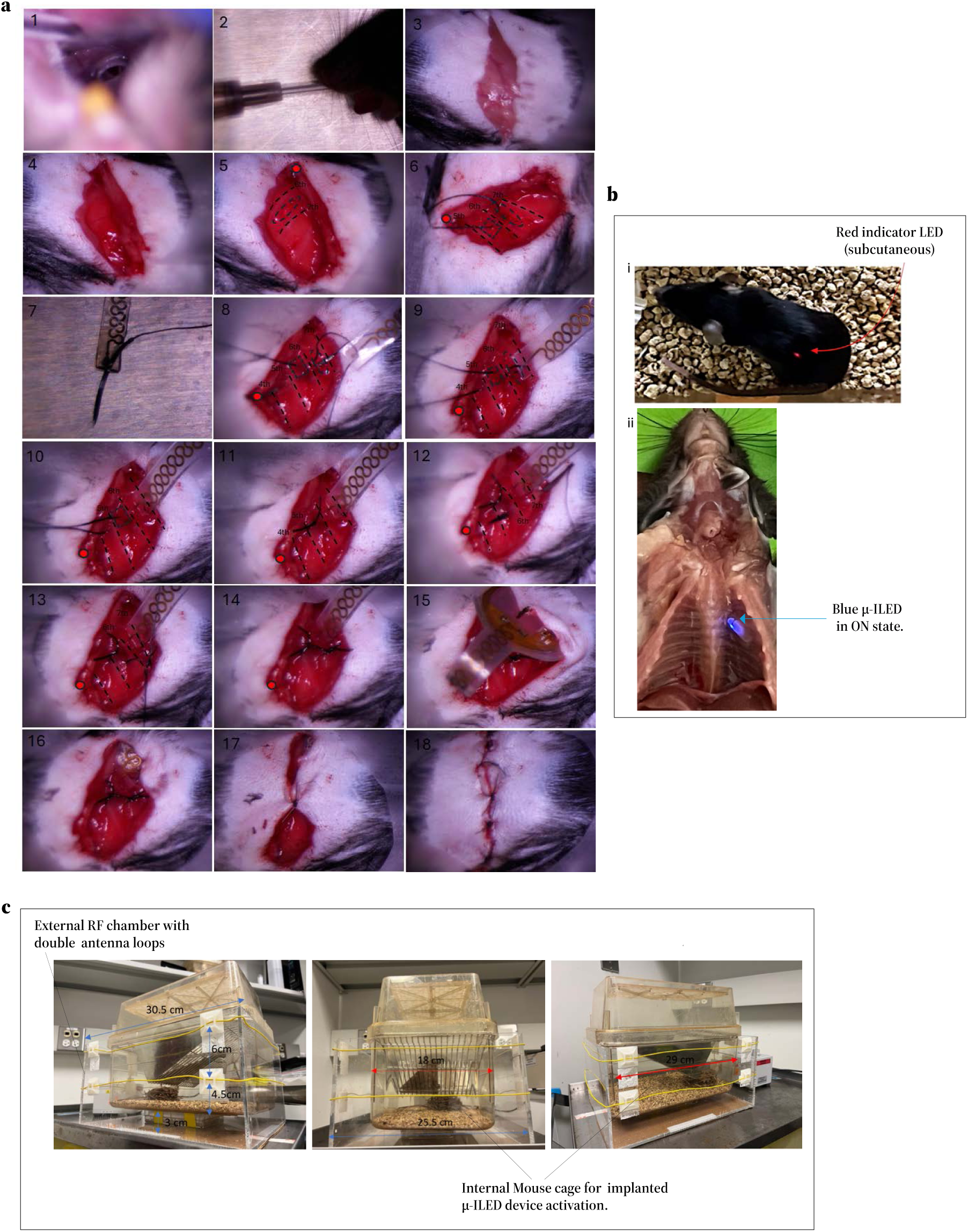
Surgical implantation of wireless optoelectronic device *in vivo*. **a**, Images of step-by-step surgical procedure for posterolateral, back-mounted, intrathoracic implantation of peripheral wireless μ-ILED device for spatiotemporal optogenetics in mouse lungs. **b**, Location of peripheral wireless device LEDs. **i**, Subcutaneous red indicator LED (650 nm) signals active wireless NFC connection; **ii**, A ventrally dissected mouse chest cavity, after removal of internal organs, showing precise location of implanted μ-ILED (470 nm) in its ON state at experimental end-point. **c**, Radio Frequency (RF) Chamber set up for wireless NFC connection.

**Supplementary Fig. 7:**
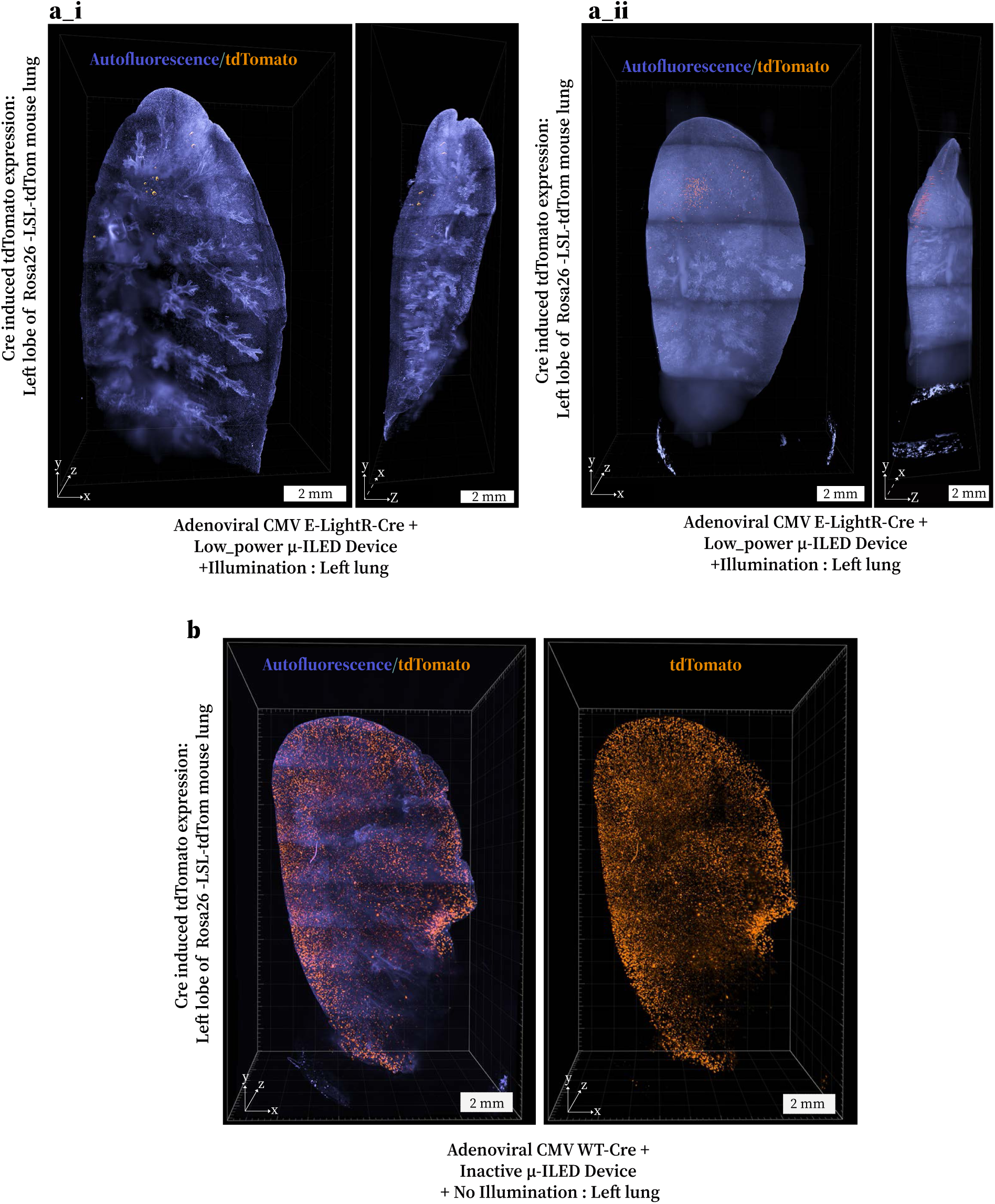
Light-mediated local genome editing in mouse lungs. **a**, **b**, Representative images of cleared left lung lobe from Rosa26-LSL tdTomato mice obtained using light sheet fluorescent microscopy (LSFM). Three-dimensional views of the left lung lobes showing reporter tdTomato expression (Orange) over background autofluorescence of lung tissue (Purple) marking the lung architecture. The recombination effects are evaluated 10 days following Cre activation. **a_i,ii**, Images from mice that received adenoviral E-LightR-Cre and local illumination with Low_power-μ-ILED device (470 nm; 10 W, 10 Hz, 50% duty cycle, 4-24 Hour). Local activation radius (95% signal area): i, ≤ 2.3 mm; ii, ≤ 3.2 mm. **b**, Images of the left lung from a mouse that received adenoviral WT-Cre. Scale bars 2 mm.

**Supplementary Fig. 8:**
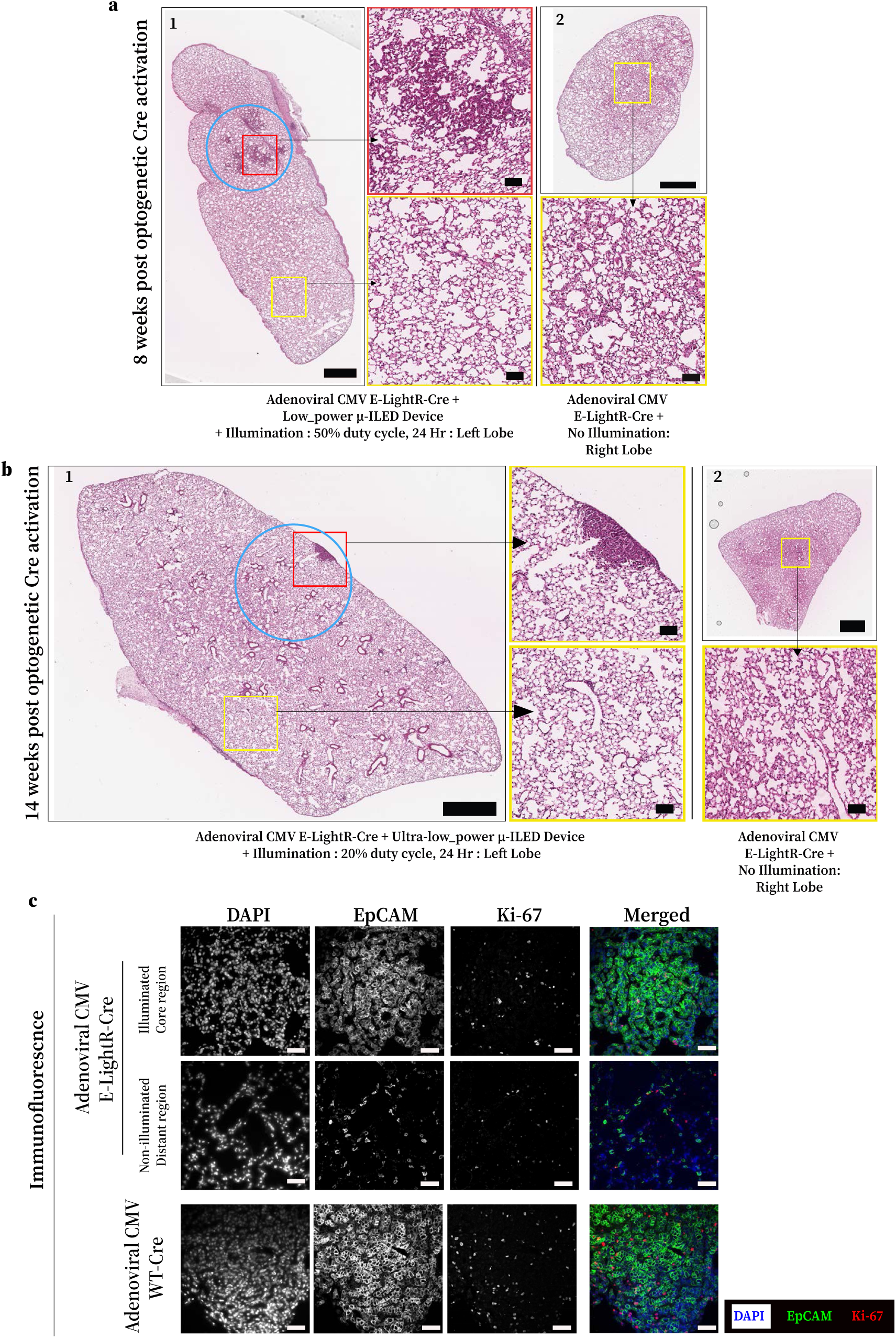
Light-mediated induction of spatially restricted tumorigenesis in mouse lungs. **a**, **b**, Representative H&E images from additional lung samples at 8-14 weeks after endotracheal adenoviral E-LightR-Cre delivery and local illumination on the left lung lobe, using μ-ILEDs. **a**, Local illumination was performed using Low_power-μ-ILED device (470 nm; 10 W, 10 Hz, 50% duty cycle, 24 Hour) and lungs were harvested 8 weeks post activation. Panel 1 shows the left lung with the illuminated area outlined by a blue circle. Zoomed-in images show illuminated (red rectangle) area and non-illuminated distant (yellow rectangle) region. Panel 2 depicts a representative right lung lobe from the same mouse with zoomed-in region (yellow rectangle). Scale bars represent 1 mm for whole-lobe images and 100 µm for zoomed-in regions. **b**, Local illumination was performed using Ultra-low_power-μ-ILED device (470 nm; 10 W, 10 Hz, 20% duty cycle, 24 Hour) and lungs were harvested 14 weeks post activation. Panel 1 shows the left lung with the illuminated area outlined by a blue circle. Zoomed-in images show illuminated (red rectangle) area and non-illuminated distant (yellow rectangle) region. Panel 2 depicts a representative right lung lobe from the same mouse with zoomed-in region (yellow rectangle). Scale bars represent 1 mm for whole-lobe images and 100 µm for zoomed-in regions. **c**, Representative images of lung sections from indicated group of mouse samples, co-stained (IF) for Ki-67 and EpCAM. Scale bars 100 μm.

