## supplemental tables for "Local optogenetic control of genome editing and tumorigenesis *in vivo* using wireless implantable optoelectronics"

### Table 1: Simulated tissue illumination volumes at varying irradiance thresholds

| Conditions | ≥ 0.05 mW/mm² | ≥ 0.1 mW/mm² | ≥ 0.25 mW/mm² | ≥ 0.5 mW/mm² | |
|---|---|---|---|---|---|
| **Standard_power** | 1.98E+08 | 1.04E+08 | 3.59E+07 | 1.17E+07 | **Volume ($\mu m^3$)** |
| **Low_power** | 3.14E+07 | 9.60E+06 | 1.91E+05 | 0 | |
| **Ultra-low_power** | 1.91E+05 | 0 | 0 | 0 | |

**Table 2: Number of genes with significant differential expression amongst the specified cell populations under study**

| Cell populations | | # Significant Differentially Expressed Genes (Padj<0.05) | # Upregulated (log2FC>0) | # Downregulated (log2FC<0) |
|---|---|---|---|---|
| | Comparison | | | |
| GFP-positive | mTmG(GFP-KRasG-12D) vs mTmG(GFP-only) (S3 vs S1) | 3798 | 1219 | 2579 |
| | mTmG(GFP-KRas-WT) vs mTmG(GFP-only) (S2 vs S1) | 59 | 14 | 45 |
| | mTmG(GFP-KRas-G12D) vs (GFP-KRas-WT) (S3 vs S2) | 2106 | 941 | 1165 |
| GFP-negative | mTmG(GFP-KRasG-12D) vs mTmG(GFP-only) (S3 vs S1) | 13 | 6 | 7 |
| | mTmG(GFP-KRas-WT) vs mTmG(GFP-only) (S2 vs S1) | 7 | 3 | 4 |
| | mTmG(GFP-KRas-G12D) vs (GFP-KRas-WT) (S3 vs S2) | 5 | 2 | 3 |

**Table 3: List of genes with significant differential expression amongst the GFP negative groups under study**

| GFP-negative: mTmG(GFP-only) vs mTmG(GFP-KRas-G12D) | | | | | | | |
|---|---|---|---|---|---|---|---|
| | Gene | BaseMean | Log2 FoldChange | Standard error value (lfcSE) | Stat | pvalue | padj |
| 1 | ENSG00000273217 | 162.0233764 | 20.23807074 | 2.330488321 | 8.684047267 | 3.8192996283115e-18 | 9.84195321219592e-14 |
| 2 | C1orf52 | 536.2915205 | 0.986171612 | 0.171748154 | 5.741963399 | 9.35850527767599e-09 | 0.00012058 |
| 3 | TBC1D3D | 57.84119572 | 7.785047627 | 1.417572586 | 5.491815872 | 3.97822031097778e-08 | 0.000341716 |
| 4 | ENSG00000290018 | 161.7401048 | -24.83879027 | 4.606741014 | -5.391835615 | 6.97415546384413e-08 | 0.000449293 |
| 5 | ZNF322P1 | 14.59256703 | -7.413487018 | 1.544656726 | -4.799439831 | 1.59110026849369e-06 | 0.008200213 |
| 6 | ENSG00000285542 | 24.45593413 | -6.635987052 | 1.421639893 | -4.667839645 | 3.04383309305644e-06 | 0.013072756 |
| 7 | ENSG00000267261 | 27.14419124 | 17.65774464 | 3.832636054 | 4.607206213 | 4.08115220732555e-06 | 0.013626426 |
| 8 | RNU6-36P | 35.526495 | -23.61914816 | 5.134895651 | -4.599732841 | 4.23033117707393e-06 | 0.013626426 |
| 9 | CRYAB | 158.2917544 | 2.467638156 | 0.549916499 | 4.48729609 | 7.21327948716242e-06 | 0.020653222 |
| 10 | CTSB | 2127.708474 | -0.710603395 | 0.166105573 | -4.278022585 | 1.88560893123789e-05 | 0.04163741 |
| 11 | ENSG00000285816 | 51.73795669 | 3.196851512 | 0.751505867 | 4.253927548 | 2.10053291195701e-05 | 0.04163741 |
| 12 | P2RX4 | 95.79023182 | -2.496422124 | 0.582614567 | -4.284860461 | 1.82853917780088e-05 | 0.04163741 |
| 13 | ZNF204P | 57.1821748 | -9.521259427 | 2.229583862 | -4.27041996 | 1.95105232506919e-05 | 0.04163741 |
| **GFP-negative: mTmG(GFP-KRas-WT) vs mTmG(GFP-KRas-G12D)** | | | | | | | |
| | Gene | BaseMean | Log2 Fold Change | Standard error value (lfcSE) | Stat | pvalue | padj |
| 1 | ENSG00000279095 | 78.20830647 | 22.60731293 | 3.665666633 | 6.167312851 | 6.9460246287652e-10 | 2.45618376897766e-05 |
| 2 | PRKAR2A-AS1 | 17.93023985 | -27.91944345 | 5.135404665 | -5.436658894 | 5.42889267394739e-08 | 0.000959855 |
| 3 | ENSG00000285542 | 24.45593413 | -7.072938274 | 1.416303906 | -4.993941092 | 5.91594492649346e-07 | 0.006973124 |
| 4 | C1orf52 | 536.2915205 | 0.822327835 | 0.170664119 | 4.818399082 | 1.44714690355122e-06 | 0.01279314 |
| 5 | IL16 | 19.9691337 | -3.973330505 | 0.869427344 | -4.570054686 | 4.8759696308756e-06 | 0.034483832 |

**Table 4 : Amino Acid sequence Information of light regulated Cre recombinase and modified mTmG reporter constructs**

| Light regulated Cre constructs | |
|---|---|
| Color Codes | **Color Codes** |
| | N'-Cre<br>SV40 NLS<br>Cre-VVD Linker<br>N'-VVD<br>Inter-VVD Linker<br>C'-VVD<br>C'-Cre<br>miRFP670<br>Y324F |
| LightR-Cre-miRFP | **LightR-Cre-miRFP670**[45] |
| | MVPKKKRKVSNLLTVHQNLPALPVDATSDEVRKNLMDMFRDRQAFSEHTWKMLLSVCRSWAA<br>WCKLNNRKWFPAEPEDVRDYLLYLQARGLAVKTIQQHLGQLNMLHRRSGLPRPSDSNAVSLVM<br>RRIRKENVDAGERAKQALAFERTDFDQVRSLMENSGPGGSGGHTLYAPGGYDIMGYLIQIMNRP<br>NPQVELGPVDTSCALILCDLKQKDTPIVYASEAFLYMTGYSNAEVLGRNCRFLQSPDGMVKPKS<br>TRKYVDSNTINTMRKAIDRNAEVQVEVVNFKKNGQRFVNFLTMIPVRDETGEYRYSMGFQCETE<br>GGSGGSGGSGGSGGGSGGSGGSHTLYAPGGYDIMGYLIQIMNRPNPQVELGPVDTSCALILCD<br>LKQKDTPIVYASEAFLYMTGYSNAEVLGRNCRFLQSPDGMVKPKSTRKYVDSNTINTMRKAIDR<br>NAEVQVEVVNFKKNGQRFVNFLTMIPVRDETGEYRYSMGFQCEGSGGPGGRCQDIRNLAFLGI<br>AYNTLLRIAEIARIRVKDISRTDGGRMLIHIGRTKTLVSTAGVEKALSLGVTKLVERWISVSGVADD<br>PNNYLFCRVRKNGVAAPSATSQLSTRALEGIFEATHRLIYGAKDDSGQRYLAWSGHSARVGAA<br>RDMARAGVSIPEIMQAGGWTNVNIVMNYIRNLDSETGAMVRLLEDGDRILQSTVPRARDPPVAT<br>MVAGHASGSPAFGTASHSNCEHEEIHLAGSIQPHGALLVVSEHDHRVIQASANAAEFLNLGSVL<br>GVPLAEIDGDLLIKILPHLDPTAEGMPVAVRCRIGNPSTEYCGLMHRPPEGGLIIELERAGPSIDLS<br>GTLAPALERIRTAGSLRALCDDTVLLFQQCTGYDRVMVYRFDEQGHGLVFSECHVPGLESYFGN<br>RYPSSTVPQMARQLYVRQRVRVLDVTYQPVPLEPRLSPLTGRDLDMSGCFLRSMSPCHLQFL<br>KDMGVRATLAVSLVVGGKLWGLVVCHHYLPRFIRFELRAICKRLAERIATRITALES- |
| E-LightR-Cre-miRFP | **E-LightR-Cre-miRFP670** |
| | MVPKKKRKVSNLLTVHQNLPALPVDATSDEVRKNLMDMFRDRQAFSEHTWKMLLSVCRSWAA<br>WCKLNNRKWFPAEPEDVRDYLLYLQARGLAVKTIQQHLGQLNMLHRRSGLPRPSDSNAVSLVM<br>RRIRKENVDAGERAKQALAFERTDFDQVRSLMENSGSGGHTLYAPGGYDIMGYLIQIMNRPNPQ<br>VELGPVDTSCALILCDLKQKDTPIVYASEAFLYMTGYSNAEVLGRNCRFLQSPDGMVKPKSTRK<br>YVDSNTINTMRKAIDRNAEVQVEVVNFKKNGQRFVNFLTMIPVRDETGEYRYSMGFQCETEGEF<br>DIRFRTDDDEQFEKVLKEMNRRARKDAGHTLYAPGGYDIMGYLIQIMNRPNPQVELGPVDTSCA<br>LILCDLKQKDTPIVYASEAFLYMTGYSNAEVLGRNCRFLQSPDGMVKPKSTRKYVDSNTINTMRK<br>AIDRNAEVQVEVVNFKKNGQRFVNFLTMIPVRDETGEYRYSMGFQCEGSGGRCQDIRNLAFLGI<br>AYNTLLRIAEIARIRVKDISRTDGGRMLIHIGRTKTLVSTAGVEKALSLGVTKLVERWISVSGVADD<br>PNNYLFCRVRKNGVAAPSATSQLSTRALEGIFEATHRLIYGAKDDSGQRYLAWSGHSARVGAA<br>RDMARAGVSIPEIMQAGGWTNVNIVMNYIRNLDSETGAMVRLLEDGDRILQSTVPRARDPPVAT<br>MVAGHASGSPAFGTASHSNCEHEEIHLAGSIQPHGALLVVSEHDHRVIQASANAAEFLNLGSVL<br>GVPLAEIDGDLLIKILPHLDPTAEGMPVAVRCRIGNPSTEYCGLMHRPPEGGLIIELERAGPSIDLS<br>GTLAPALERIRTAGSLRALCDDTVLLFQQCTGYDRVMVYRFDEQGHGLVFSECHVPGLESYFGN<br>RYPSSTVPQMARQLYVRQRVRVLDVTYQPVPLEPRLSPLTGRDLDMSGCFLRSMSPCHLQFL<br>KDMGVRATLAVSLVVGGKLWGLVVCHHYLPRFIRFELRAICKRLAERIATRITALES- |

| | |
|---|---|
| CI-LightR-Cre-miRFP | **CI-LightR-Cre-miRFP670** |
| | MVPKKKRKVSNLLTVHQNLPALPVDATSDEVRKNLMDMFRDRQAFSEHTWKMLLSVCRSWAA<br>WCKLNNRKWFPAEPEDVRDYLLYLQARGLAVKTIQQHLGQLNMLHRRSGLPRPSDSNAVSLVM<br>RRIRKENVDAGERAKQALAFERTDFDQVRSLMENSGPGGSGGHTLYAPGGYDIMGYLIQIMNRP<br>NPQVELGPVDTSCALILCDLKQKDTPIVYASEAFLYMTGYSNAEVLGRNCRFLQSPDGMVKPKS<br>TRKYVDSNTINTMRKAIDRNAEVQVEVVNFKKNGQRFVNFLTMIPVRDETGEYRYSMGFQCETE<br>GGSGGSGGSGGSGGGSGGSGGSHTLYAPGGYDIMGYLIQIMNRPNPQVELGPVDTSCALILCD<br>LKQKDTPIVYASEAFLYMTGYSNAEVLGRNCRFLQSPDGMVKPKSTRKYVDSNTINTMRKAIDR<br>NAEVQVEVVNFKKNGQRFVNFLTMIPVRDETGEYRYSMGFQCEGSGGPGGRCQDIRNLAFLGI<br>AYNTLLRIAEIARIRVKDISRTDGGRMLIHIGRTKTLVSTAGVEKALSLGVTKLVERWISVSGVADD<br>PNNYLFCRVRKNGVAAPSATSQLSTRALEGIFEATHRLIYGAKDDSGQRYLAWSGHSARVGAA<br>RDMARAGVSIPEIMQAGGWTNVNIVMNFIRNLDSETGAMVRLLEDGDRILQSTVPRARDPPVAT<br>MVAGHASGSPAFGTASHSNCEHEEIHLAGSIQPHGALLVVSEHDHRVIQASANAAEFLNLGSVL<br>GVPLAEIDGDLLIKILPHLDPTAEGMPVAVRCRIGNPSTEYCGLMHRPPEGGLIIELERAGPSIDLS<br>GTLAPALERIRTAGSLRALCDDTVLLFQQCTGYDRVMVYRFDEQGHGLVFSECHVPGLESYFGN<br>RYPSSTVPQMARQLYVRQRVRVLVDVTYQPVPLEPRLSPLTGRDLDMSGCFLRSMSPCHLQFL<br>KDMGVRATLAVSLVVGGKLWGLVVCHHYLPRFIRFELRAICKRLAERIATRITALES- |
| **Reporter Constructs** | |
| mTmG[2] | **Original Reporter GFP sequence in mTmG after Cre recombination** |
| | VSKGEELFTGVVPILVELDGDVNGHKFSVSGEGEGDATYGKLTLKFICTTGKLPVPWPTLVTTLT<br>YGVQCFSRYPDHMKQHDFFKSAMPEGYVQERTIFFKDDGNYKTRAEVKFEGDTLVNRIELKGID<br>FKEDGNILGHKLEYNYNSHNVYIMADKQKNGIKVNFKIRHNIEDGSVQLADHYQQNTPIGDGPVL<br>LPDNHYLSTQSALSKDPNEKRDHMVLLEFVTAAGITLGMDELYK- |
| mTmG(kRAS-G12D) | **Reporter GFP-KRas-G12D sequence in mTmG(GFP-KRas-G12D) after Cre recombination** |
| | VSKGEELFTGVVPILVELDGDVNGHKFSVSGEGEGDATYGKLTLKFICTTGKLPVPWPTLVTTLT<br>YGVQCFSRYPDHMKQHDFFKSAMPEGYVQERTIFFKDDGNYKTRAEVKFEGDTLVNRIELKGID<br>FKEDGNILGHKLEYNYNSHNVYIMADKQKNGIKVNFKIRHNIEDGSVQLADHYQQNTPIGDGPVL<br>LPDNHYLSTQSALSKDPNEKRDHMVLLEFVTAAGITLGMDELYKSLGGPSGSTMTEYKLVVVGA<br>DGVGKSALTIQLIQNHFVDEYDPTIEDSYRKQVVIDGETCLLDILDTAGQEEYSAMRDQYMRTGE<br>GFLCVFAINNTKSFEDIHHYREQIKRVKDSEDVPMVLVGNKCDLPSRTVDTKQAQDLARSYGIPFI<br>ETSAKTRQGVDDAFYTLVREIRKHKEKMSKDGKKKKKKSKTKCVIM- |
| mTmG(kRAS-WT) | **Reporter GFP-KRas-WT sequence in mTmG(GFP-KRas-WT) after Cre recombination** |
| | VSKGEELFTGVVPILVELDGDVNGHKFSVSGEGEGDATYGKLTLKFICTTGKLPVPWPTLVTTLT<br>YGVQCFSRYPDHMKQHDFFKSAMPEGYVQERTIFFKDDGNYKTRAEVKFEGDTLVNRIELKGID<br>FKEDGNILGHKLEYNYNSHNVYIMADKQKNGIKVNFKIRHNIEDGSVQLADHYQQNTPIGDGPVL<br>LPDNHYLSTQSALSKDPNEKRDHMVLLEFVTAAGITLGMDELYKSLGGPSGSTMTEYKLVVVGA<br>GGVGKSALTIQLIQNHFVDEYDPTIEDSYRKQVVIDGETCLLDILDTAGQEEYSAMRDQYMRTGE<br>GFLCVFAINNTKSFEDIHHYREQIKRVKDSEDVPMVLVGNKCDLPSRTVDTKQAQDLARSYGIPFI<br>ETSAKTRQGVDDAFYTLVREIRKHKEKMSKDGKKKKKKSKTKCVIM- |

**Table 5: List of plasmids and adenovirus**

| Plasmid List | | | |
|---|---|---|---|
| **Construct name** | **Description** | **Source** | **Used in Figure** |
| LightR-Cre miRFP670[45] | CMV promoter, pN1 backbone | (Addgene plasmid # 162158 ; http://n2t.net/addgene:162158 ; RRID:Addgene_162158). | Fig. 1, Suppl. Fig. 1 |
| CI-LightR-Cre miRFP670 | CMV promoter, pN1 backbone, Y324F mutation in Original Cre recombinase sequence[29] | Made in the Lab; Site directed mutagenesis. | Fig. 1, Suppl. Fig. 1 |
| E-LightR-Cre miRFP670 | CMV promoter, pN1 backbone | Made in the Lab; Modified Site directed Mutagenesis[49]. | Fig. 1, 3 and Suppl. Fig. 1, 5 |
| Lv-SD-PA-Cre-nMag_Opti[30] | CAG Promoter, Lentiviral Backbone | A gift from Dr. Masayuki Yazawa. | |
| LiCre(AslOv2)[34] | CMV promoter; AsLOV2-CreE340AD341A; PGAE0 Self-Inactivating Vector | pGY577 was a gift from Dr. Gaël Yvert (Addgene plasmid # 166663; http://n2t.net/addgene:166663; RRID:Addgene_166663). | |
| PA Cre 3.0 miRFP670 | CMV promoter, pN1 backbone | Made in the Lab; generation of megaprimer followed by modified site directed mutagenesis as previously described[49] . | Fig. 1 and Suppl. Fig. 1 |
| LiCre(AsLOV2) miRFP670 | CMV promoter, pN1 backbone | Made in the Lab; generation of megaprimer followed by modified site directed mutagenesis[49]. | Fig. 1 and Suppl. Fig. 1 |
| pCA-mTmG[46] | pCA-HZ2 Backbone | PCA-mTmG was a gift from Dr. Liqun Luo (Addgene plasmid # 26123; http://n2t.net/addgene:26123; RRID:Addgene_26123). | Fig. 1, 3 and Suppl. Fig. 1, 5 |
| Floxed-STOP-mCherry[29] | pcDNA3.1 backbone | pcDNA3.1_Floxed-STOP mCherry was a gift from Moritoshi Sato (Addgene plasmid # 122963; http://n2t.net/addgene:122963; RRID:Addgene_122963). | Fig. 3 |

**Table 5 continued**

| Construct name | Description | Source | Used in Figure |
|---|---|---|---|
| pBabe-Kras WT | pBabe Backbone, Retroviral Backbone | pBabe-Kras Wt was a gift from Dr. Channing Der (Addgene plasmid # 75282 ; http://n2t.net/addgene:75282 ; RRID:Addgene_75282). | |
| pBabe-KRas G12D | pBabe Backbone, Retroviral Backbone | pBabe-Kras G12D was a gift from Dr. Channing Der (Addgene plasmid # 58902 ; http://n2t.net/addgene:58902 ; RRID:Addgene_58902). | |
| pCA-mTmG(GFP-KRas -WT) | pCA-HZ2 Backbone | Made in the Lab; generation of megaprimer followed by modified site directed mutagenesis[49]. | Fig. 3 and Supplementary Fig. 5 |
| pCA-mTmG(GFP-KRas-G12D) | pCA-HZ2 Backbone | Made in the Lab; generation of megaprimer followed by modified site directed mutagenesis[49]. | Fig. 3 and Supplementary Fig. 5 |
| pShuttle[84] | Mammalian Expression, Adenoviral | pShuttle was a gift from Dr. Bert Vogelstein (Addgene plasmid # 16402 ; http://n2t.net/addgene:16402 ; RRID:Addgene_16402). | |
| **Adenovirus list** | | | |
| **Adenovirus Name** | **Description** | **Source** | **Used in Figure** |
| Adenoviral WT-Cre-eGFP | Adenovirus. CMV promoter driven. | A gift from Dr. Kyle Schachtschneider. | Fig. 5 and Suppl. Fig. 7, 8 |
| Adenoviral E-LightR-Cre-miRFP | Adenovirus. CMV promoter driven. | Synthesized at RRC Viral Vector Core, UIC, Chicago, IL. | Fig. 5 and Suppl. Fig. 7, 8 |

**Supplementary File S1: Detailed protocol for surgical implantation of peripheral wireless optoelectronic device *in vivo*.**

**Equipment list**

- Stereo Microscope (AmScope, SM-3T Series Trinocular LED; cat. no SM-3TZ-54S-5M)
- Cotton tipped applicators (ULINE, cat. no S-21102)
- Hair clipper (Wahl, cat. no 79434)
- Heating equipped Small Animal surgery platform (Harvard Apparatus, cat. no 50-1239)
- Heating equipped Surgical recovery Platform (Kent Scientific, cat. no SURGI-M)
- Glass bead sterilizer (Cole-Parmer, cat. no UX-10779-00)
- Anesthesia vaporizer (Drager Vapor 19.1 isoflurane)
- Intubation stand (Kent Scientific, cat. no ETI-MSE-01)
- Ventilator (Harvard Apparatus, cat. no 55-7040)
- Sterile surgical air (Linde Gas & Equipment, cat. no AI M-K)
- Insulin syringe (Exel Int., cat. no 26027)
- 21G needle (BD Surgical, cat. no 305167)
- 1mL syringe (BD Surgical, cat. no 309659)
- 20G x 1 ¼” catheter (Exel Int., cat no. 26742)
- Hartman Mosquito hemostatic forceps (Kent Scientific, cat. no INS750452)
- Vessel clip (WPI, cat. no 15911)
- Micro-spatula, flat rounded/tapered (Millipore Sigma, cat. no Z513334)
- Reverse cutting P-3 needle (13 mm, 3/8) with 6-0 silk sutures (AD Surgical, cat. no S-S618R13)
- Needle holder (Kent Scientific, cat. no INS600109)
- McPherson-Vannas scissors (Kent Scientific, cat. no INS600124)
- Iris forceps (Kent Scientific, cat. no INS650915)
- Fine dressing forceps (Kent Scientific, cat. no 650914)
- Reverse cutting P-3 needle (13 mm, 3/8) with 6-0 Nylon sutures (AD Surgical, cat. no S-N610R13-B)
- wireless, battery-free, miniaturized, fully-implantable optoelectronic devices containing blue (470 nm) $\mu$-ILED.

**Reagents list**

- Systane lubricant eye ointment (Alcon, cat. no 1635970-0623)
- Depilatory cream (Nair)
- Ethanol 70%
- Ketamine/xylazine
- Isoflurane (Covertus, cat. no 29405)
- Buprenorphine ER (Wedgewood Connect, cat. no 79926-058-17)

**Preoperative preparation**

1. Sterilize all surgical instruments by autoclaving. If surgery is performed on multiple mice within the same session, the instruments can be re-sterilized between mice by removing

all visible material with 70% ethanol solution and then sterilizing with a glass bead sterilizer.

2. Anesthetize the mouse using 100/5 mg/kg body weight ketamine/xylazine via intraperitoneal injection.
3. Once anesthetized completely, deliver Buprenorphine ER (1mg/ml) via a subcutaneous injection between the shoulder blades of the mice.
4. Shave the surgical site with a hair clipper.
5. Place the mouse onto the intubation stand, suspending the mouse from its maxillary incisors and securing the limbs with tape.
6. Intubate the mouse with a 20G catheter (Supplementary Fig. 6a_1).

   **Critical Step:** Visualize the proper space to insert the catheter for intubation using the stereo microscope. Use a vessel clip held with a hemostatic forceps to hold the tongue out of the way, further aided with a micro-spatula to properly visualize the trachea for catheter placement. Improper placement will prevent proper ventilation during the procedure, increasing the risk of harm to the animal.
7. Remove the mouse from the intubation stand and place in a right lateral recumbent position on the heating plate positioned under the dissecting microscope. Secure limbs and tail with tape.
8. Connect the mouse to the ventilator and begin flow of sterile surgical air with 1% isoflurane (Supplementary Fig. 6a_2).
9. Apply eye lubricant to the eyes to prevent drying of the cornea during surgery.
10. Apply Nair to the previously shaved surgical area to ensure all hair is removed from the skin.

    **Critical Step:** Take care to remove all hair from the surgical site. If any hair remains at the site of incision, it will contaminate the surgical field.
11. Scrub the skin with 70% ethanol.

**Surgical implantation procedure**

12. Make the first cut approximately over the 5$^{th}$ rib, approximately 1cm in length, through the skin layer, using McPherson-Vannas scissors (Supplementary Fig. 6a_3).
13. Cut through all muscle layers as necessary and spread them apart until the rib bones are visualized and the left lobe of the lung is visible within the chest cavity (Supplementary Fig. 6a_4).
14. Using the McPherson-Vannas scissors, make an incision into the chest cavity between the 6$^{th}$ and 7$^{th}$ rib approximately the size of the tip of the NeuroLux peripheral device (3mm) (Supplementary Fig. 6a_5). This incision should be approximately 10mm from the spinal cord laterally.
15. Rotate the entire heating plate 90 degrees so the mouse in now vertical, head facing upwards. Insert a reverse cutting P-3 needle (13 mm, 3/8) with a 6-0 silk suture through the 5th and 6th rib, staying closer to the 5th rib, and exiting the thoracic cavity through the incision space between the 6th and 7th ribs, made in step 14. (Supplementary Fig. 6a_6). Return the heating plate to the original position.

    **Critical Step:** Take precautions to avoid any needle-contact to the lung and any risk of causing damage to the lung thereby, while inserting and withdrawing the suture within the chest cavity.

16. Insert the suture needle into the tip of the NeuroLux device next to the μ-ILED, inserting into the top of the device and exiting through the bottom (Supplementary Fig. 6a_7).

    **Critical Step:** Ensure the device is in the proper orientation. The receiver contains a red LED indicator that should be facing upwards as that should be how it is inserted into the subcutaneous space later in procedure step 24.

17. Insert the suture needle back through the incision made in step 14 to reenter the chest cavity, then exit through the intercostal muscle between the 4$^{th}$ and 5$^{th}$ rib, staying close to the 5$^{th}$ rib (Supplementary Fig. 6a_8).
18. Pull both ends of the suture to move the NeuroLux device into position, to be inserted into the chest cavity (Supplementary Fig. 6a_9).

    **Critical Step:** Make one final check to ensure device is in the proper orientation with the indicator LED facing upwards.

19. Pull both ends of the suture and guide the tip of the NeuroLux device into the chest cavity with iris forceps until the μ-ILED is fully underneath the ribs, inside the chest cavity (Supplementary Fig. 6a_10).
20. Tie the suture securely around the 5$^{th}$ rib (Supplementary Fig. 6a_11).
21. Again, using the 6-0 silk suture, seal the chest cavity by first preparing to tie a suture around the 6$^{th}$ and 7$^{th}$ ribs. Enter the chest cavity through the 7$^{th}$ intercostal space and exiting through the 5$^{th}$ intercostal space directly laterally to the insertion incision of the NeuroLux device. Loosely tighten this suture, but do not tie off (Supplementary Fig. 6a_12).
22. Perform an inspiration hold on the ventilator for 4 seconds. During this time, gently squeeze the chest cavity to remove any excess air and pull the suture tight (Supplementary Fig. 6a_13).

    **Critical Step:** Sealing the chest cavity is a requirement to reestablish the negative pressure system by which the lungs operate. Removing the excess air and forming a proper seal is required for the mouse to survive the surgery.

23. Finish tying the suture to completely seal the chest cavity (Supplementary Fig. 6a_14).
24. Create a subcutaneous cavity in the dorsal region of the animal using fine dressing forceps to prepare for inserting the wireless signal receiver end of the NeuroLux device (Supplementary Fig. 6a_15).
25. Insert the device into the created cavity to position the receiver end so it will be parallel to the ground with the red LED facing upwards when the animal is upright (Supplementary Fig. 6a_16).

    **Critical Step:** Proper insertion of the serpentine coil and the receiver across the body of the mouse will lead to proper activation ability using the NeuroLux optogenetic system. Take care to ensure the serpentine coil is lying flat with no twists or bends so that it will keep the receiver in the proper position and will minimize movement after surgery.

26. Close the surgical incision using interrupted 6-0 nylon skin sutures (Supplementary Fig. 6a_17,18).

**Postoperative care**

27. At the conclusion of the surgical procedure, stop the flow of isoflurane but keep the mouse on the ventilator until there are clear signs of the mouse beginning to breathe on its own.

28. Once proper breathing has been established by the mouse without the need of the ventilator, extubate.
29. Transfer the mouse to the recovery heating platform and give supplemental flow of sterile air while monitoring the mouse closely until it begins to show signs of recovery from anesthesia.
30. Return the mouse to the housing cage and allow 7-10 days of recovery until performing adenovirus delivery and optical stimulation.

**Supplementary File S2: Detailed protocol for analyzing animal locomotor behavior following wireless optoelectronic device implantation.**

1. Image J with Animal Tracker Plugin load: [AnimalTracker - Download (elte.hu)]
2. Video capture: 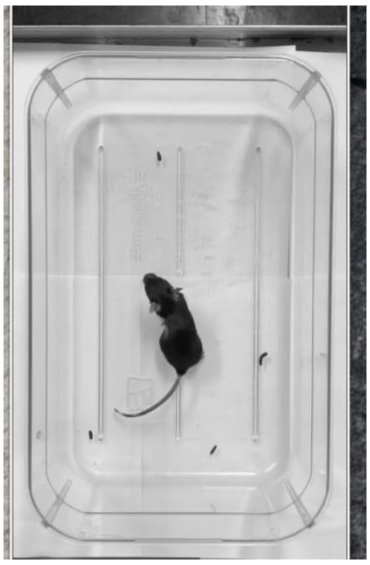
   Capture for ~30 minutes.
   Note: The videos were taken with iphone 13 at HD settings with 25 frames per second (fps) and 2.5X magnification. The Cage bottom is 11.2 inches (28.448 cm) long.
3. Convert videos to Image Sequence: use Free studio
4. Open Image J
5. File > Import > Image Sequence >

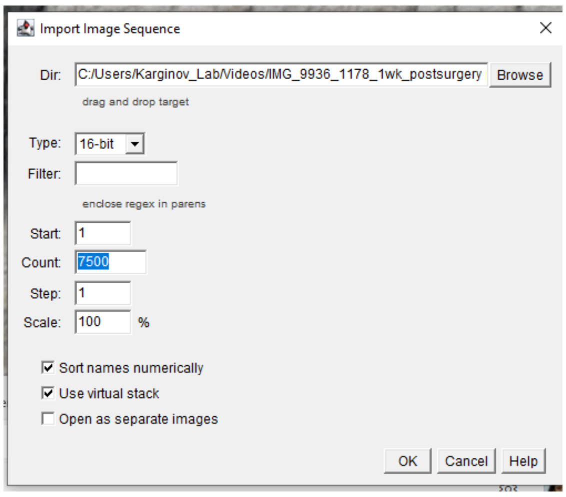


6. Click on Image sequence > on Image J menu, select polygon selection > make a polygon around area of interest, in this case a rectangular shape around the cage area.

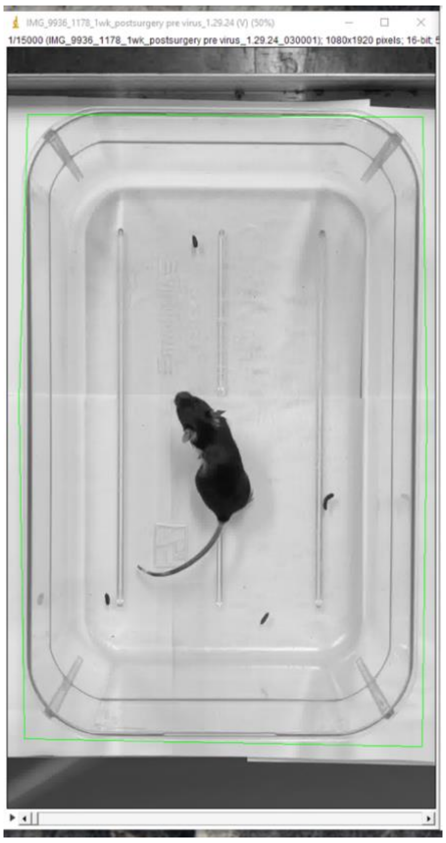

7. Plugins> Animal Tracker > examples > Radial Maze
8. Click on image sequence, click set image> set maze> show tracker.
9. Click on filter from drop down box on the Processing window.

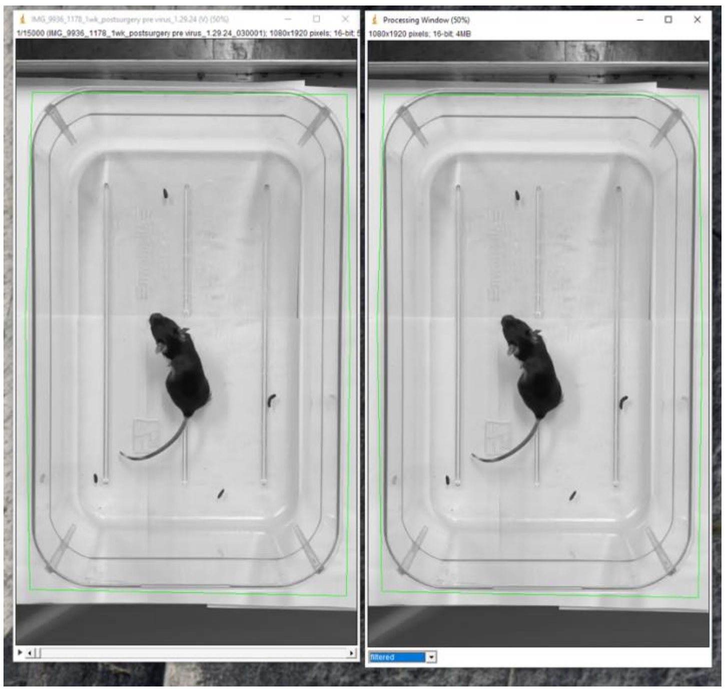

10. A Tracker window will also appear on screen.
11. Set original image sequence as active window.
12. Set filters>  Background Subtraction > set image sequence as active image > add 3-4 frames with nonoverlapping mouse location > click on show filter >

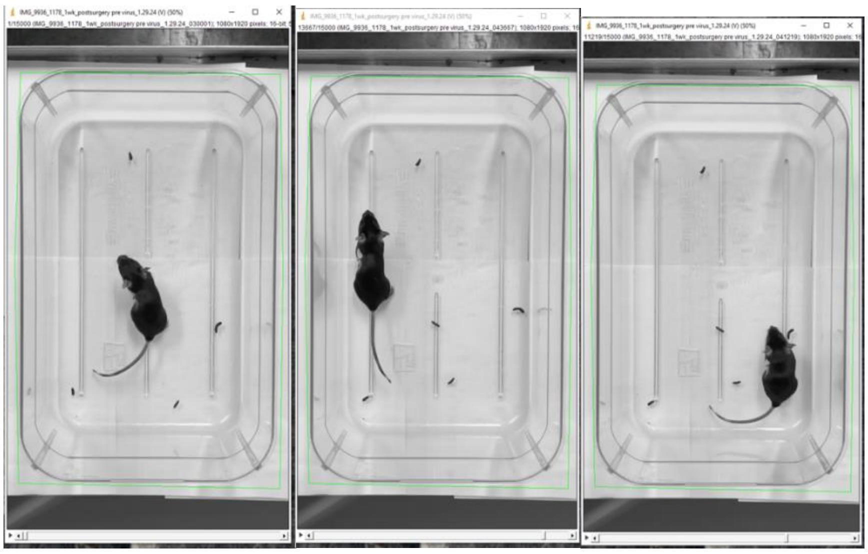

Note: Ideal filter background subtraction will look like below. With no mouse in cage.

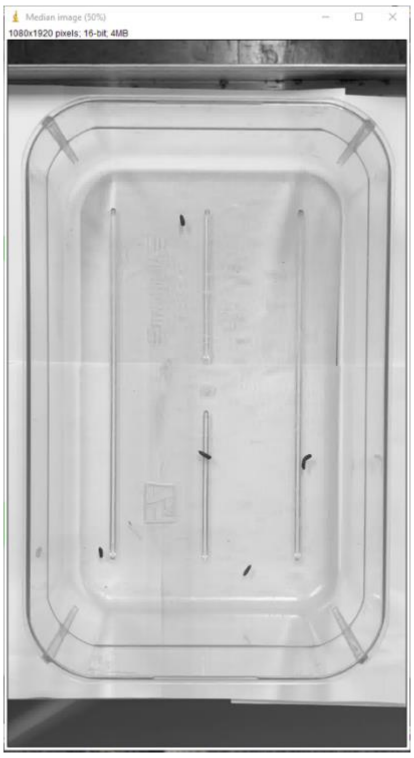

13. Click Done. Cross out the median image.
14. Go to the Filter Settings > Select Gaussian Blur > Add > set at 50
15. Click Ok to get out of Filter settings
16. Drag image sequence in original image sequence to see the filtered image in Processing window.

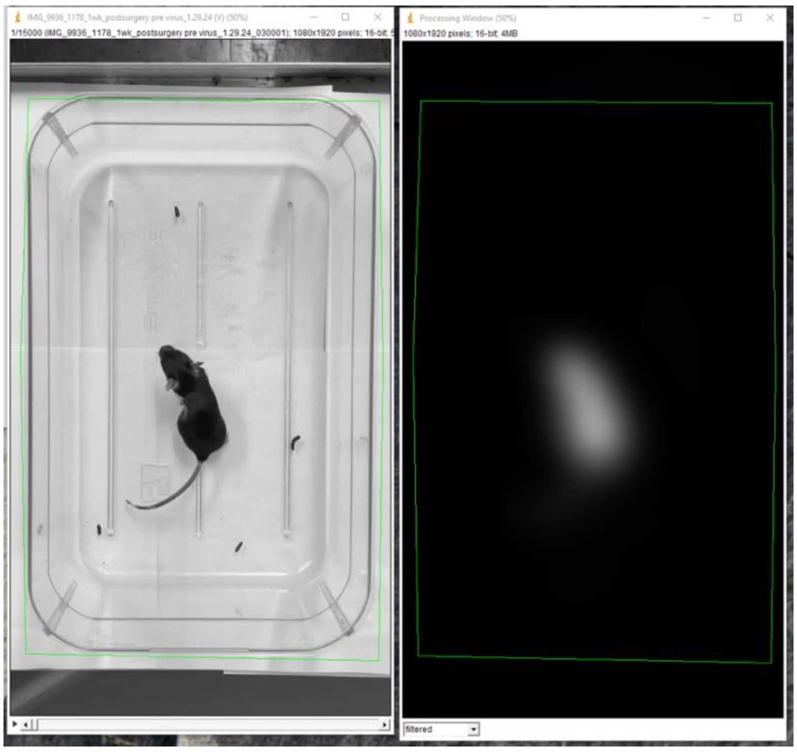

17. Click processing window> set threshold > click grey thresholder >adjust threshold on mouse body > click ok

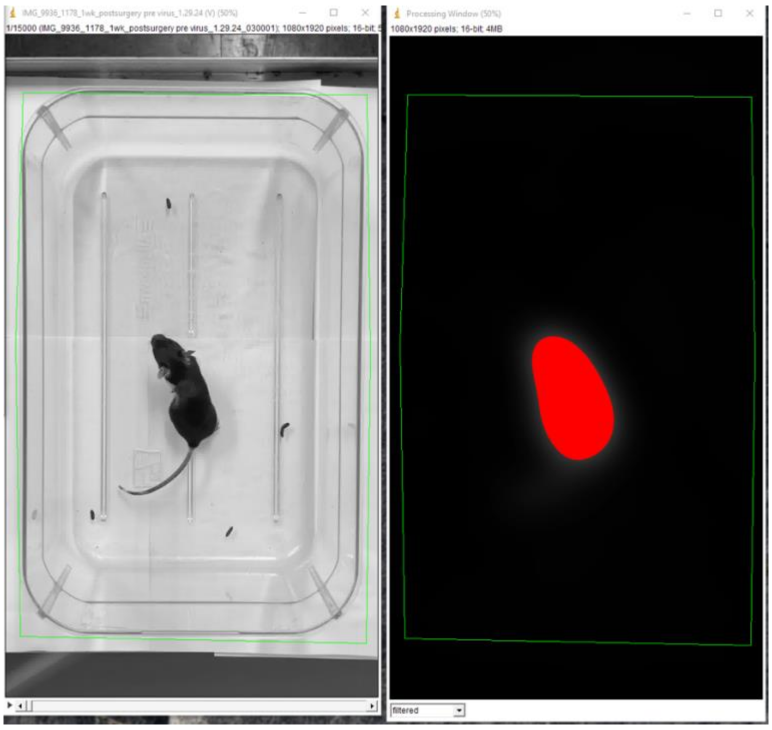

18. Click on original image sequence> On tracker: set last frame > 7500/15000
19. Click on original image sequence> On tracker: show blobs: Yellow box will show up around thresholded area over mouse. > double click on the blob on main image sequence, it will turn read.
20. Click Strat Tracking on Tracker menu.

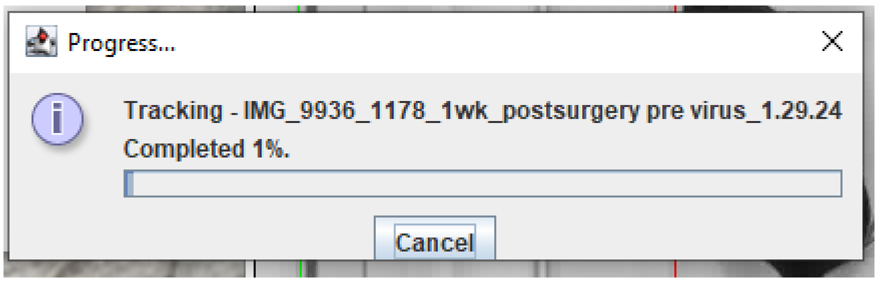


21. Track finished: Click Ok.

22. Click on Show analyser. Select “All” Zones

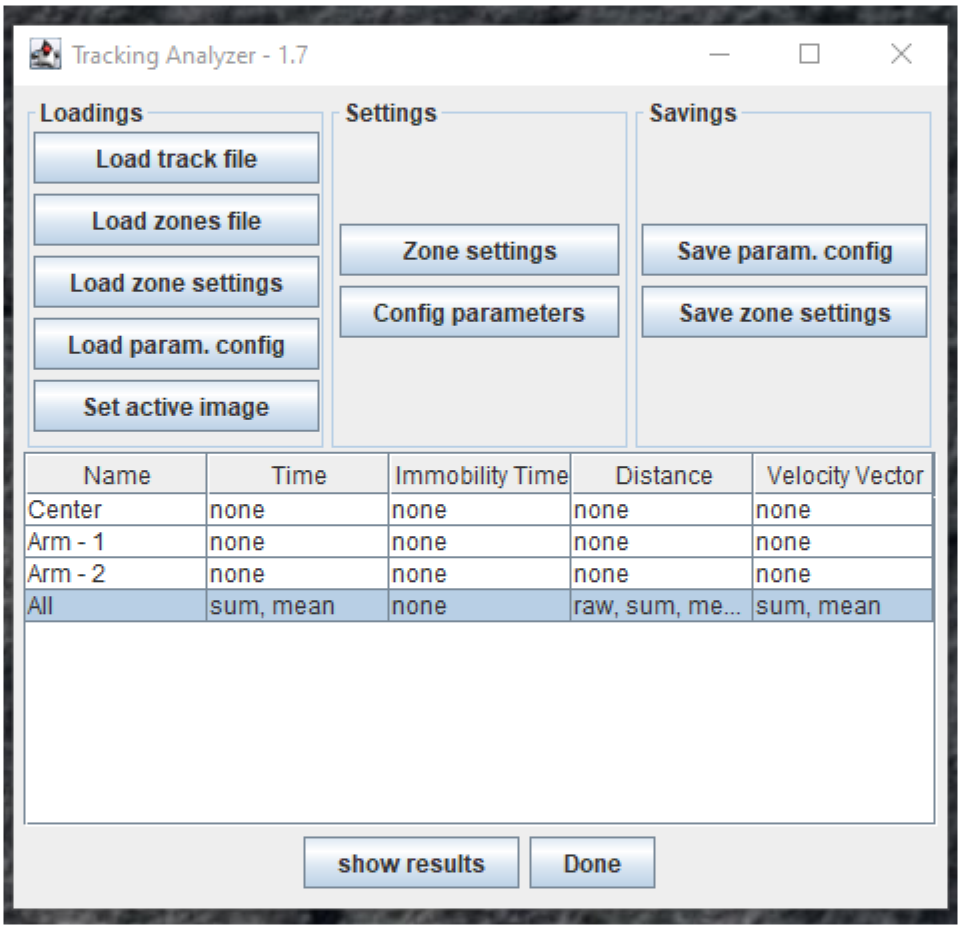


23. Go to Zone settings> Set Parameters>

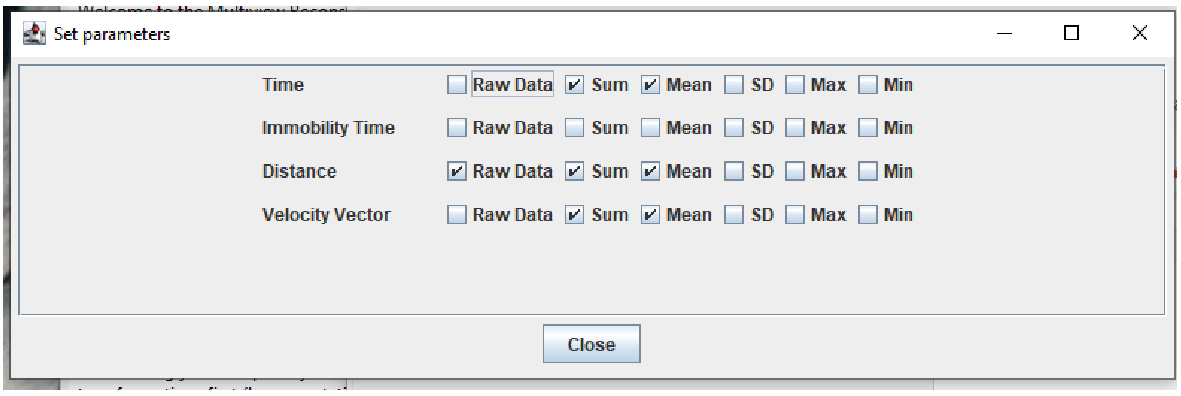


24. For fps 25; time settings are as follows:

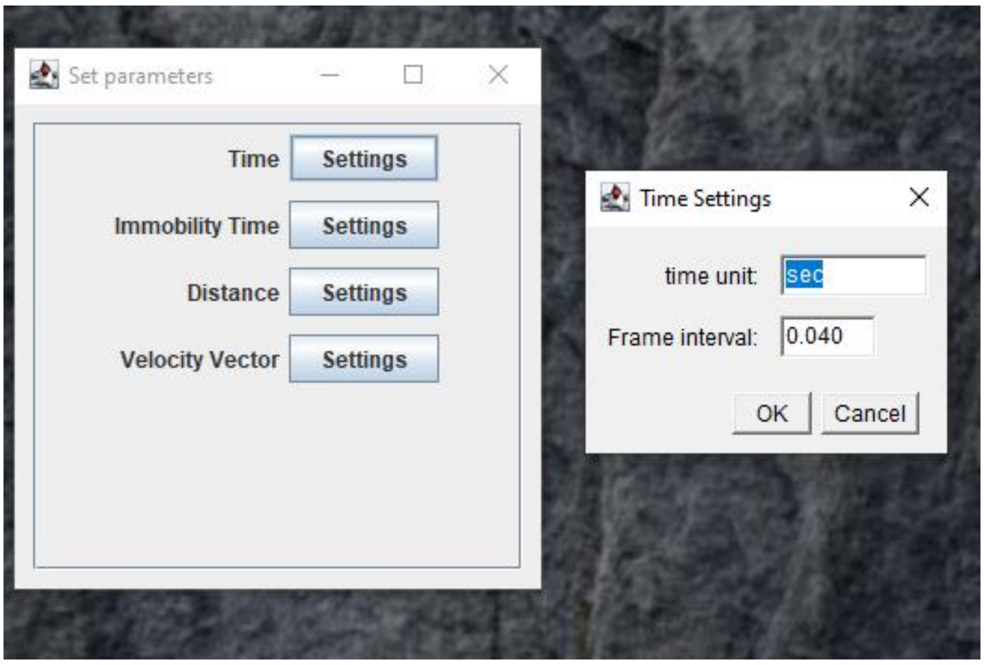


25. Distance settings: cage pixel length / cage length in cm

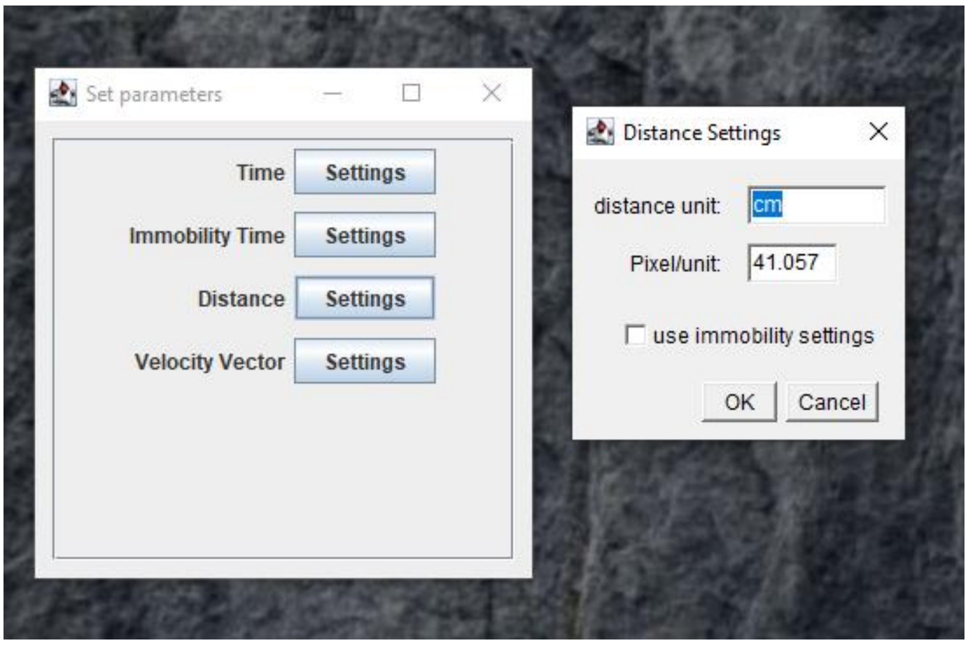


26. Velocity settings:

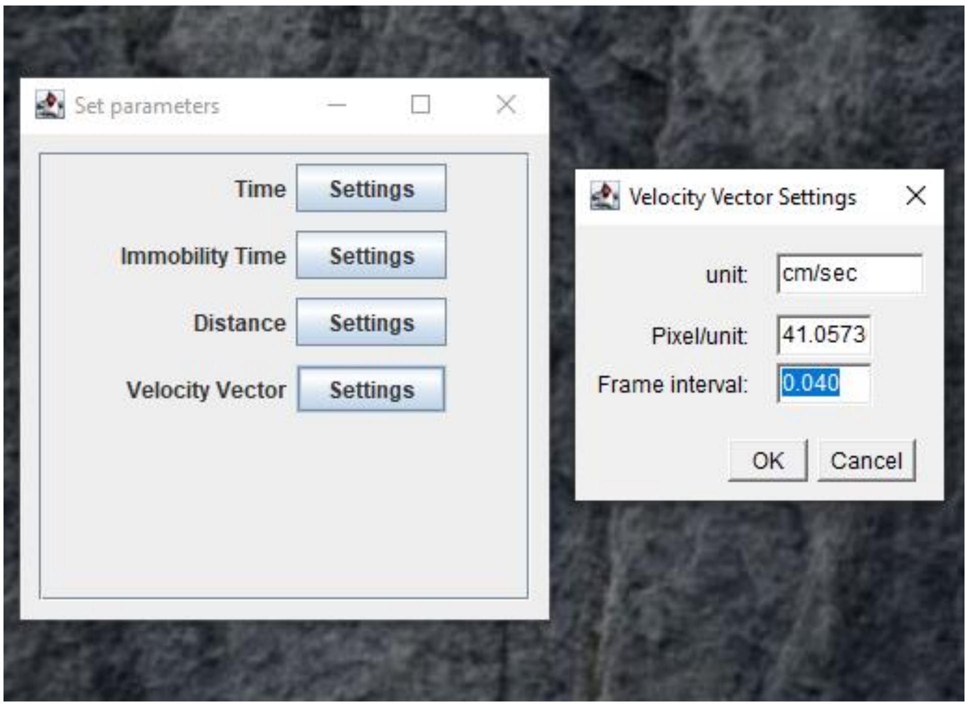


27. Click all zones on list > Click Show Results > Grouped by Zones > Copy and Paste result window on Excell Sheet
28. Use data of every 5 minutes to measure:
    A. Avg distance travelled in 5 minutes from n=3 of 5 minute intervals from each video.
    B. Avg Velocity in 5 minutes from n=3 of 5 minute intervals from each video.

Note: The images and values shown in the screenshots are illustrative and intended solely for method description; they do not represent results from specific experiments.
